# *Etv6/Runx1* Fusion Gene Abrogation Decreases The Oncogenic Potencial Of Tumour Cells In A Preclinical Model Of Acute Lymphoblastic Leukaemia

**DOI:** 10.1101/809525

**Authors:** Adrián Montaño, Jose Luis Ordoñez, Verónica Alonso-Pérez, Jesús Hernández-Sánchez, Teresa González, Rocío Benito, Ignacio García-Tuñón, Jesús María Hernández-Rivas

**Author notes:** These authors share senior authorship. **Corresponding author:** Jesús-María Hernández-Rivas, IBMCC, CIC Universidad de Salamanca-CSIC, Hospital Universitario de Salamanca, Paseo de San Vicente 58, 37007 Salamanca, Spain, //.

## Abstract

**Background:** The t(12;21)(p13;q22), which fuses *ETV6* and *RUNX1* genes, is the most common genetic abnormality in children with B-cell precursor acute lymphoblastic leukaemia. The implication of the fusion protein in leukaemogenesis seems to be clear. However, its role in the maintenance of the disease continues to be controversial.

**Aim:** To eliminate the expression of the *ETV6/RUNX1* fusion gene, in order to elucidate the effect in the leukaemic cells.

**Methods:** Generation of an *in vitro ETV6/RUNX1* knock out model using the genetic modification system CRISPR/Cas9. Functional studies and generation of edited-cell xenograft model were carried out.

**Results:** For the first time, the expression of ETV6/RUNX1 fusion gene was completely eliminated, thus generating a powerful model on which to study the role of the fusion gene in leukaemic cells. ETV6/RUNX1 inactivation caused the deregulation of cellular processes that could be participating in the maintenance of the leukaemic phenotype, such as differentiation and lymphoid activation, apoptosis, cell signaling and cell migration. Tumour cells showed higher levels of apoptosis, lower proliferation rate and a greater sensitivity to PI3K inhibitors *in vitro* along as a decrease in tumour growth in xenografts models after *ETV6/RUNX1* fusion gene abrogation.

**Conclusions:** ETV6/RUNX1 fusion protein plays a fundamental role in the maintenance of the leukaemic phenotype, thereby being making the fusion protein a potential therapeutic target.

## BACKGROUND

The gene fusion between the transcription factors *ETV6* (*TEL*) and *RUNX1* (*AML1*) is generated by t(12;21)(p13; q22), the most frequent chromosomal translocation in children with acute lymphoblastic leukaemia (ALL).^1,2^ Patients carrying this translocation are associated with a good prognosis and excellent molecular response to treatment. However up to 20% of cases relapse.^3–7^ Furthermore, the response to treatment of some relapse cases is associated with resistance to treatments such as glucocorticoids (GCs),^8^ and these patients must be treated with stem cell transplantation.^9^

ETV6/RUNX1 (E/R) protein is known to play a role in the development of B-ALL, but by itself it is not able of initiating the disease. Postnatal genetic events are required for the development of clinically overt leukaemia. These second events are usually mutations or deletions, such as the loss of wild type (WT) allele of *ETV6.*^10^ Recent studies suggest that E/R is responsible for the initiation of leukaemia and also essential for disease progression and maintenance, through deregulation of different molecular pathways that contribute to leukaemogenesis. E/R regulates phosphoinositide 3-kinase (PI3K)/Akt/mammalian target of rapamycin (mTOR) (PI3K/AKT/mTOR) pathway, which promotes proliferation, cell adhesion and DNA damage response; *STAT3* pathway involved in self-renewal and cell survival and *MDM2/TP53* whose deregulation induces the inhibition of apoptosis and consequently cell survival.^11^ Therefore, the fusion protein could thus be intervening in relapse processes and in the lack of response to treatment.

However, the functional studies carried out by the silencing of *E/R* fusion gene expression, mediated by siRNA and shRNA, reveal that there is still controversy about the role of the oncoprotein in the maintenance of the leukaemic phenotype. Thus E/R silencing by siRNA neither induced cell cycle arrest/apoptosis nor attenuated clonogenic potential of cells. Therefore the E/R fusion protein may be dispensable for the survival of definitive leukaemic cells.^12^ By contrast, other studies showed that E/R expression was critical for the survival and propagation of the respective leukaemia cells in *vitro* and in *vivo.*^13^ These results arise some doubts about the implications of the fusion protein in tumour cells.

The implementation of new genetic editing strategies has allowed the development of functional studies by generation of gene and gene fusion KO models, both *in vitro* and *in vivo.*^14^ In this study, we completely abrogated the expression of E/R fusion protein in REH ALL cell line using the CRISPR/Cas9 editing system and we studied the effects of genetic ablation of the fusion protein in cellular functions. We also studied whether the suppression of E/R expression sensitize tumour cells to PI3K inhibitors. Finally, we generated a xenograft mouse model to study the oncogenic potential of tumours cells after E/R depletion. In summary, we provide evidence that fusion protein has a key role in the maintenance of the leukaemic phenotype.

## MATERIAL AND METHODS

### Cell lines and culture conditions

REH, obtained from DMSZ German collection (ACC 22), is a cell line established from the peripheral blood of a patient with ALL who carried t(12;21) and del(12) producing respective *E/R* fusion and deletion of residual *ETV6*. REH was maintained in RPMI 1640 (Life Technologies, Carlsbad, California, USA) supplemented with 15% fetal bovine serum (FBS) and 1% of Penicillin/ Streptomycin (P/S) (Life Technologies). Stromal HS-5 cell line was obtained from ATCC collection (CRL-11882) and maintained in DMEM (Life Technologies, Carlsbad, California, USA) supplemented with 10% FBS and 1% of P/S. Both cells lines were maintained at 37ºC with 5% CO2.

### sgRNAs design and cloning

Based on the methodology of CRISPR/Cas9, two single guides RNAs (sgRNAs) (G1 and G2) were designed with the Broad Institute CRISPR designs software (http://www.broadinstitute.org/rnai/public/analysis-tools/sgrna-design). One of them directed towards the end of exon 5 of *ETV6* and other directed towards the beginning of intron 5-6, both before the fusion point, with the intention of producing indels or deletions that modify the open reading frame of the oncogene, and, therefore, the gene expression. These sgRNAs were cloned into a vector containing the Cas9 nuclease coding sequence and GFP, pSpCas9(BB)-2A-GFP (PX458) (Addgene plasmid #48138)^15^ as described previously^14^ (Supplementary Table 1). Then they were electroporated into the REH cells.

### sgRNA transfections

REH ALL cells (4 × 10 ^6^ cells) were electroporated with 4 µg of both plasmid constructs ^14^ (PX458 G1 and PX458 G2) using the Amaxa electroporation system (Amaxa Biosystem, Gaithersburg, MD, USA) according to supplier’s protocol.

### Flow cytometry analysis and cell sorting

Seventy-two hours after sgRNAs transfection, GFP-positive cells were selected by fluorescence-activated cell sorting (FACS) using FACS Aria (BD Biosciences, San Jose, California, USA). Single-cells were seeded in 96-well plate by FACS, establishing the different KO and control clones.

### Sequencing of sgRNA targets sites

Genomic DNA was extracted using the QIAamp DNA Micro Kit (Qiagen, Hilden, Germany) following the manufacturer’s protocol. To amplify the region of *E/R* fusion, PCR was performed using the following primers: forward 5′-ACCCTCTGATCCTGAACCCC– 3′ and reverse 5′-GGATTTAGCCTCATCCAAGCAG– 3′. PCR products were purified using a High Pure PCR Product Purification Kit (Roche, Basilea, Switzerland) and were sequenced by the Sanger method using each forward and reverse PCR primers (Supplementary table 2).

The editing efficiency of the sgRNAs and the potential induced mutations were assessed using Tracking of Indels by Decomposition (TIDE) software (https://tide-calculator.nki.nl; Netherlands Cancer Institute), which only required two Sanger sequencing runs from wild-type cells and mutated cells.

### Off-target sequence analysis

The top four predicted off-target sites obtained from “Breaking Cas” website (http://bioinfogp.cnb.csic.es/tools/breakingcas/) were analyzed by PCR in the different clones (Supplementary table 2) before to functional and xenograft experiments.

### qPCR

Total RNA extraction was performed with the RNeasy Kit (Qiagen) as suggested by the manufacturer. Real-time reverse transcriptase– polymerase chain reactions (RT-PCRs) were performed as described ^16^. The primers for *E/R* (sense, 5 - CTCTGTCTCCCCGCCTGAA −3; antisense, 5 - CGGCTCGTGCTGGCAT-3), were designed. Real-time RT-PCR data shown include at least 3 independent experiments with 3 replicates per experiment.

### Transcriptome sequencing

RNA-seq was performed by using SMART-Seq v4 Ultra Low Input RNA kit (Clonthech, California, U.S.). In all samples, RNA was analyzed following manufacturer’s recommendations for the protocol “Illumina library preparation using Covaris shearing”. Libraries were sequenced in the HiSeq400 platform (Illumina) according to manufacturer’s description with a read length of 2□×□150 nucleotides. Briefly, bcl files were demultiplexing on BaseSpace (Illumina Cloud based resource) to generate fastq files. Raw data quality control was performed with fastQc (v0.11.8), globin contamination was assessed with HTSeq Count, FastQ screen evaluated ribosomal RNA contamination and other external possible resources of contamination (Mus musculus, Drosophila melanogaster, Caenorhabditis elegans and mycoplasma). STAR (v020201) was used for the alignment (hg19 reference genome) and FeatureCounts (v1.4.6) to generate the read count matrix. Finally, DESeq2 was used for differentially gene expression analysis. Of noted, DESeq2 model internally corrects for library size therefore normalizes the values and enables paired comparisons. Heatmap was performed in R.

Go enrichment analysis (http://geneontology.org) to evaluate whether a set of genes was significantly enriched between the different comparisons was used. The most significant biological mechanisms, pathways and functional categories in the data sets of genes selected by statistical analysis were identified through PANTHER Overrepresentation Test.

### Western blotting

Protein expression was assessed by SDS-PAGE and western blotting (WB). The antibodies were obtained from Cell Signaling Technology (Danvers, MA, USA) including a human anti-Bcl-2 antibody (1:1000; 2872) for Bcl-2, a human anti-Bcl-xL antibody (1:1000; 2762) for Bcl-xL, a human anti-phospho Akt antibody (1:1000; 4060) for p-AKT (Ser473) and a human anti-phospho mTOR antibody (1:1000;2971) for p-mTOR (Ser2448). Anti-rabbit IgG horseradish peroxidase-conjugated (1:5000; 7074) was used as a secondary antibody. Antibodies were detected using ECL^TM^ WB Detection Reagents (RPN2209, GE Healthcare, Illinois, Chicago, USA). ImageJ software was used for densitometric analysis.^17,18^

### Apoptosis, cell cycle analysis and proliferation assays

Apoptosis was measured by flow cytometry with an annexin V-Dy634 apoptosis detection kit (ANXVVKDY, Immunostep, Salamanca, Spain) following the manufacturer’s instructions. Briefly, 5 × 10^5^ cells were collected and washed twice in PBS and labeled with annexin V-DY-634 and non-vital dye propidium iodide (PI), allowing the discrimination of living-intact cells (annexin-negative, PI-negative), early apoptotic cells (annexin-positive, PI-negative) and late apoptotic or necrotic cells (annexin-positive, PI-positive). In parallel, cell distribution in the cell cycle phase was also analyzed by measuring DNA content (PI labeling after cell permeabilization). These experiments were carried out after 24, 48 and 72 culture hours.

For proliferation measuring, MTT assays and labeling of cells with CellTrace CFSE Cell Proliferation Kit (Thermo Fisher) were used. In MTT assays, cells were plated on 96-well plates, cell density varied according to the days of the experiment, in a range between 3×10^4^ and 5×10^3^ cells (24-240 hours). MTT solution (3-(4,5-cimethylthiazol-2-yl)-2,5-diphenyl tetrazolium bromide) was added at concentration of 0.5μg/μL (Merck, Darmstadt, Germany). After incubation for 3-4 hours at 37°C, cells were lysed with the solubilization solution (10% SDS in 0.01M HCl) and absorbance was measured in a plate reader at 570 nm. For labeling of cells, 3□×□10^5^ cells were stained with CellTrace-CFSE following the manufacturer’s instructions and plated on 6-well plates. After 48 hours, CFSE expression was measured by flow cytometry.

### B-ALL-stromal cell co-culture

HS-5 human mesenchymal stromal cells (MSCs) were plated at a density of 1□×□10^5^ cells per coverslip in 6-well plates. After 24□h, control cells and E/R KO clones were stained with Celltrace-CFSE and 3□×□10^5^ cells placed on top of the stromal cell monolayer. Cells were co-cultured during 48h in RPMI 1640 supplemented with 15% FBS and 1% of P/S at 37ºC with 5% CO2.

### Drugs and treatments

The followings drugs were used: Vincristine and Copanlisib (BAY 80-6946) were obtained from Selleckchem (Houston, USA) and Prednisolone (P6004) obtained from Merck. All drugs were prepared at the appropriate stocking concentrations in DMSO (Merck) and stored at −20ºC until use.

### Mouse xenograft tumourigenesis

16 four-to five-week-old female NOD/SCID/IL2 receptor gamma chain null (NSG) mice (Charles River, Barcelona, Spain) were used. 5 x10^6^ tumour cells from REH or control clone were subcutaneously injected into the left flank and tumour cells from KO clones (KO1, KO2 and KO3) were injected in the right flank as described previously.^14^ REH vs KO2 in the group 1, REH vs KO3 in the group 2, control clone vs KO1 in the group 3 and control clone vs KO2 in the group 4 (4 mice per group). The study received prior approval from the Bioethics Committee of our institution and followed the Spanish and European Union guidelines for animal experimentation (RD 53/2013 and 2010/63/UE).

Tumour diameters were measured every 2–3 days with a caliper. Tumour volume was calculated as described elsewhere by the formula a2bπ/6 (a and b being, respectively, the smallest and the biggest diameters). Mice were sacrificed by anesthesia overdose when tumour volume reached 2 cm^3^ or 48-62 days after cell injection, upon which the tumours were collected and weighted.

### Histopathology and immunohistochemistry

Excised tumours were sampled just after sacrifice and representative areas were a) formalin-fixed (24 hours) (Merck Millipore) and paraffin-embedded and (b) snap-frozen in OCT and stored at 80ºC as previously described.^19^ Tissue sections 2μM thick were stained with hematoxilin & eosin (H&E) and prepared for immunohistochemistry (IHC). IHC was performed as previously described^19^ using the anti-Ki67primary antibody (Merck Millipore). The number of mitotic figures were counted in 6 high-power field.

### Statistical analysis

Statistical analysis was performed using GraphPad Prism 6 Software. Differences in relative expression of E/R and cell viability after treatments were tested by Tukey’s multiple comparisons test. Differences in protein expression, proliferation and apoptosis levels were tested by unpaired *t*-test. Differences in tumour masses over time were tested by non-parametric Mann Whitney U test followed by Tukey’s multiple comparisons test and parametric Student’s t test. Statistical significance at values of *P*≤0,05 (*), *P*≤0.005 (**) and *P*≤0,001 (***) was noted.

## RESULTS

### CRISPR/Cas9 edited lymphoid cell line showed a loss of E/R functionality

*E/R* sequence was edited by CRISPR/Cas9 and evaluated by Sanger sequencing in REH cells (Supplementary Figure 1). The edition efficiency evaluated through TIDE was 62,6% with sgRNA G1 and 70,6% with sgRNA G2. The most frequent generated mutations were insertions up to 4 base pairs (bps).

Single-edited cells were seeded into a 96 well plate to obtain clones with a single edition that predicted a KO sequence for the oncogene. Only 48 single cell clones proliferated in culture. These clones were screened by sanger sequencing and the results revealed that more than 50% of clones (25/48) presented an edited *E/R* sequence. Among them, three single-edited cell clones “KO1, KO2 and KO3 clones” with a predicted *E/R* KO sequence were selected for the study. These clones had different editions in their sequences. KO1 clone carried an insertion of two cytosines at the end of exon 5, near the PAM sequence of the sgRNA G2, thus a frameshift mutation that generated a stop codon before finishing the exon. On the other hand, KO2 and KO3 had an insertion of 5 and 3 nucleotides respectively near the PAM sequence within exon 5, followed by a deletion of 100 bps approximately between both sgRNAs. These alterations modified the open reading frame, generating the stop codon in the next exon. In addition, the loss of the splicing region prevented the correct processing of the protein. Additionally, two single-edited cell clones with WT *E/R* sequence were used as control clones “Control 1 and Control 2” (Supplementary Figure 2).

In order to check the functionality of the *E/R* alleles carrying these clones, the expression of the fusion transcript E/R was quantified by quantitative PCR. For that, RNAs of the different clones were extracted and transcribed to cDNA. Quantification revealed a total loss of E/R mRNA expression in KO2 and KO3 clones as compared to control clones (*P*<0.001) and a leaky expression in KO1 clone (*P*<0.001) (Figure 1).

**Figure 1.**
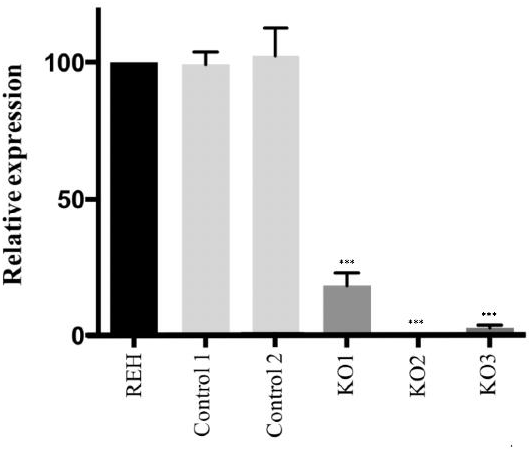
E/R expression levels by RT-qPCR. Control clones showed an expression of E/R similar to it was observed in the parental REH cells. In E/R KO clones, whose sequence was edited by the CRISPR/Cas9 system, KO2 and KO3 showed a total loss of E/R expression and KO1 showed a leaky expression. All the experiments were carried out by triplicate, the means with the standard deviations for each clone were represented. \*\*\**P≤*0.001 (unpaired *t*-test).

The four most likely off-target sequences from both guides were analyzed by sequencing Sanger into the different clones. The study of the obtained sequences revealed the lack of editing in those regions, confirming the absence of CRISPR/Cas9 off-targets (data not shown).

### Transcriptomic analysis of E/R KO lymphoid cell line generated by CRISPR/Cas9 showed a distinct expression signature and a deregulation of its downstream signaling genes

The gene expression profile analysed by total RNA-sequencing showed a distinct expression signature in E/R KO clones as compared with REH cells and control clones. 342 genes were significantly deregulated after *E/R* fusion gene depletion (q<0.05), 180 upregulated and 160 downregulated (Supplementary table 3). Some of these genes have been previously associated with ALL pathology and have also been shown to be directly regulated by *RUNX1* transcript factor^20^ (Table 1). The gene expression profile of the top50 of the most deregulated genes according to fold change (FC) values is shown in Figure 2.

**Table 1.**
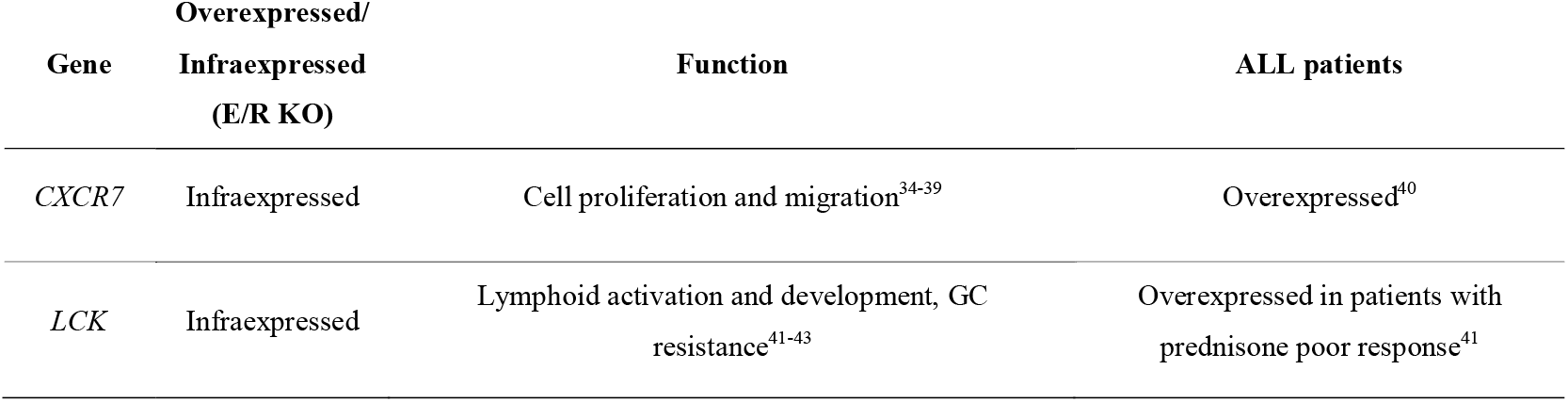
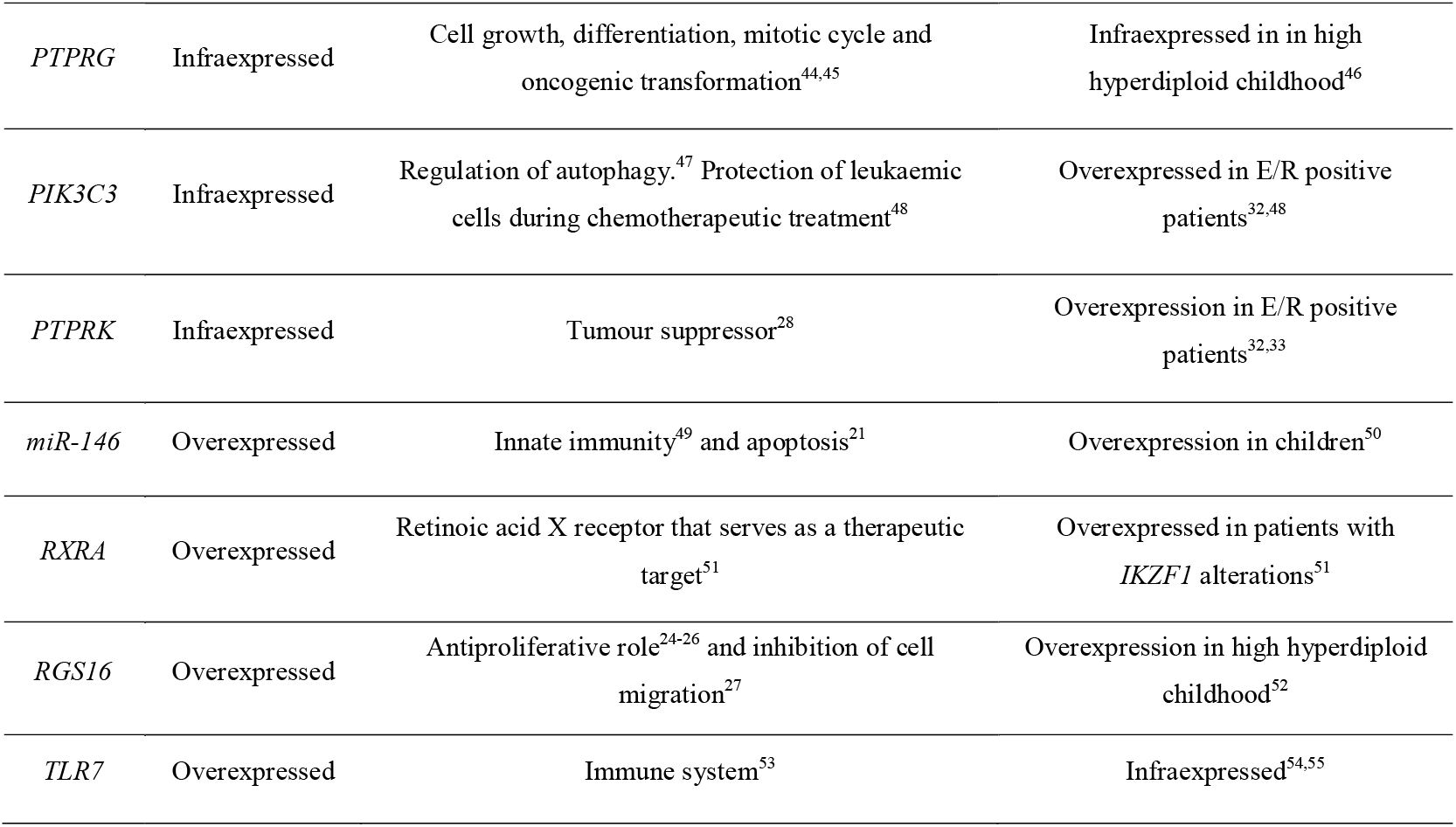
Most relevant deregulated genes associated with ALL pathology. The different columns show the name of deregulated gene, if it was overexpressed or infraexpressed after *E/R* abrogation, the cellular processes in which it participates and the expression of this gene in ALL patients.

**Figure 2.**
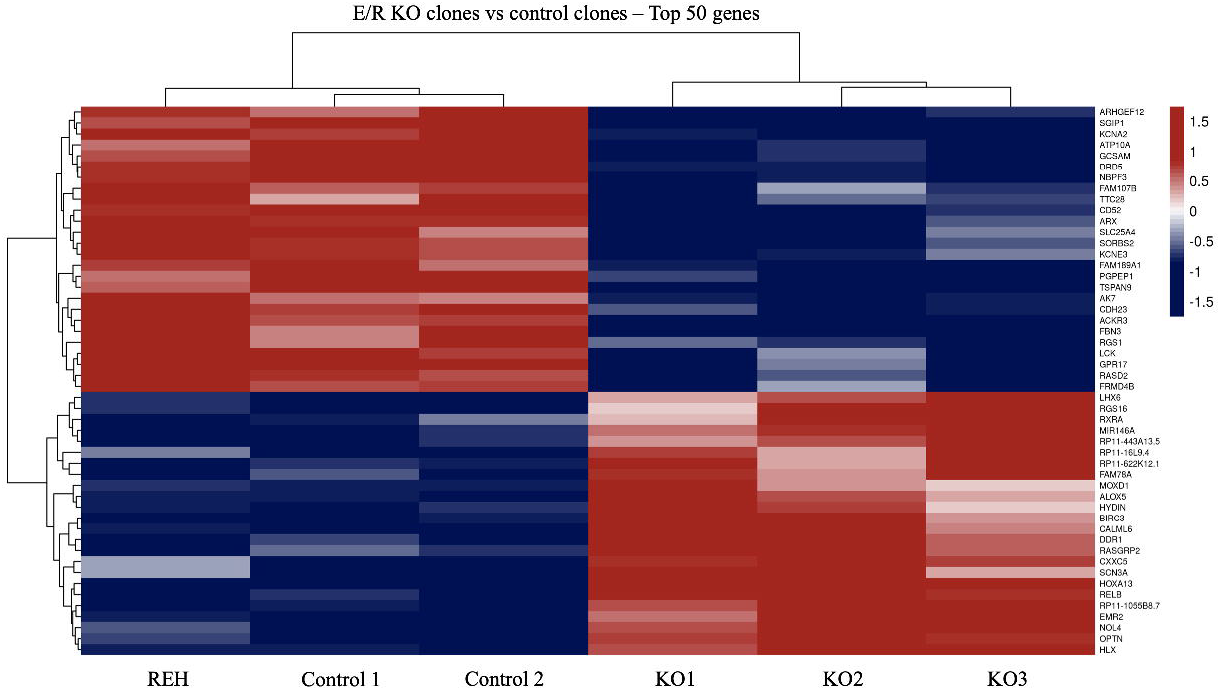
Transcriptomic analysis of E/R KO clones. Heat map of TOP50 differentially expressed genes in E/R KO clones as compared with REH cells and control clones. Each row represents one differentially expressed gene; each column represents one clone. The dendrogram on the top reveals the sample clustering; the dendrogram on the left reveals the gene clustering.

In order to elucidate the effect of *E/R* fusion gene abrogation on a functional level, the significantly deregulated genes were grouped into a cellular processes according to their function by enrichment analysis (Supplementary table 4). These cellular processes were manually classified into 11 categories: differentiation and lymphoid activation, immune response, response to stimuli, cell activation, apoptosis, cell signaling, chemical homeostasis, GPTase regulation, cell migration, location of cellular components and neuronal system development.

Apoptosis was one of the altered cellular processes. In particular, an overexpression of *miR-146* was observed in the E/R KO clones as compared with control clones (*P*<0.001). *miR-146* can regulate the expression of the apoptosis factor *STAT1*, and the anti-apoptosis factor *Bcl-xL*, thus promoting the apoptosis of ALL cells.^21^ Furthermore, tumour protein P63 gene (*TP63* gene) was also upregulated in E/R KO clones as compared with control clones (*P*<0.001). This gene is involved in an antiapoptotic pathway that regulates the normal survival of B cells.^22,23^

An overexpression of *RGS16* was also observed in ALL cells after *E/R* abrogation. This gene plays an antiproliferative role through inhibition of the PI3K/AKT/mTOR pathway^24–26^ and inhibition of cell migration.^27^ On the other hand, the tumour suppressor *PTPKR* whose expression prevents the activation of pathways such as PI3K/AKT/mTOR and STAT signaling pathways^28^ was downregulated after *E/R* fusion gene abrogation. This gene was found downregulated in some types of cancer such as ovarian, breast and NK-T cell lymphoma,^29–31^ but also in ALL cell lines.^28^ However, E/R positive ALL patients have shown an overexpression of *PTPKR.*^32,33^

### *E/R* abrogation reduces proliferative capacity and resistance to apoptosis *in vitro*

To elucidate the biological effects of abrogation of E/R expression in the KO clones, several studies were performed. MTT proliferation studies were performed at 24-hour intervals up to 240 hours. The results showed no proliferation differences between KO clones and REH cells or control clones (Supplementary Figure 3A). We simultaneously analyzed the cell cycle distribution of the different cells by permeabilization followed by PI staining. No differences were observed between the different clones (Figure 3A). No significant differences were observed through the expression of CFSE by flow cytometry (Figure 3B). However, E/R KO clones showed a significantly lower proliferation rate than control clones when they were co-cultured with MSCs (HS-5) (*P*<0.05) (Figure 3C).

**Figure 3.**
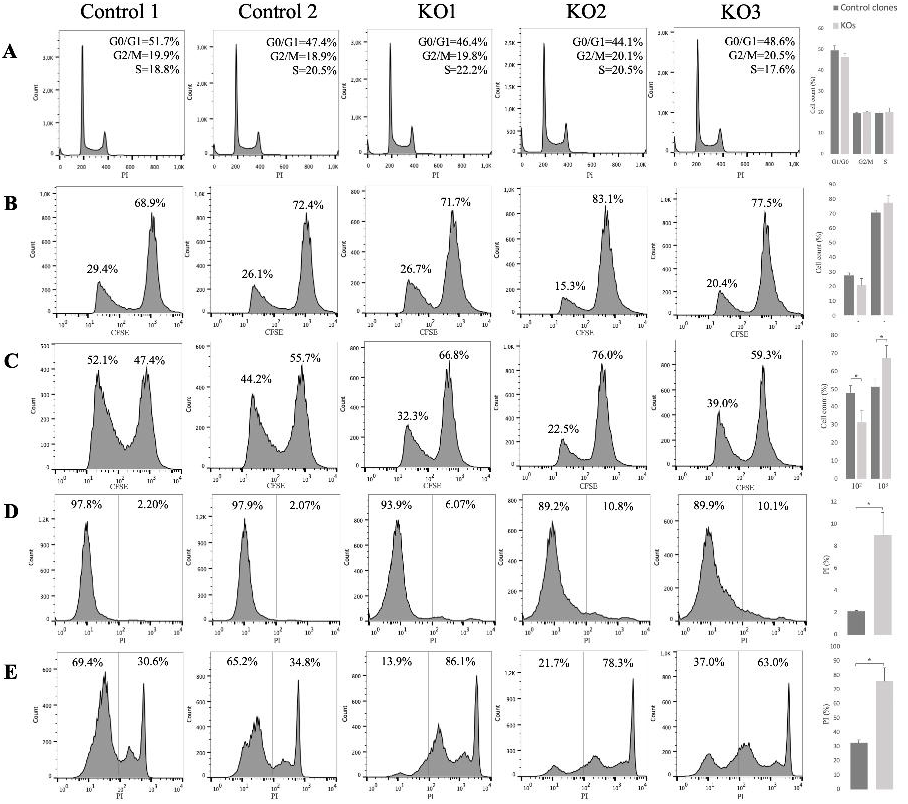
*In vitro* functional studies after E/R abrogation. (A) Cell cycle distribution of control clones and E/R KO cells at 48 h. (B) CFSE quantification by flow cytometry after 48 in culture. The peak on the right (10^3^) represents the percentage of cells that have not divided and the left peak (10^2^) represents the percentage of cells that have divided and therefore diluted their CFSE expression. (C) CFSE expression by flow cytometry of cells co-cultured with MSC cell line HS-5 at 48h. (D) Apoptosis level quantification by PI expression. The figure shows the percentage of PI negative cells (left) and PI positive cells (right) at 48 h. (E) Apoptosis level quantification by PI expression after treatment with Vincristine (1 µM) at 48 h. On the right is represented the mean distribution of control clones (dark grey) and E/R KO clones (grey) of different experiments. All the experiments were carried out by triplicate. \**P≤*0.05 (unpaired *t*-test).

Deregulation of genes such as *miR-146a* or *TP63* observed by expression analysis suggested the alteration of cellular processes such as the regulation of apoptosis. Levels of anti-apoptotic factor such us *Bcl-2* or *Bcl-xL* gene have shown to play a key role in the survival of E/R positive cells, protecting from programmed death.^56,57^ To check these findings, Bcl-2 and Bcl-xL expression levels were measured through WB. Suppression of the fusion protein produced a decrease of 60% and 47% in the expression of Bcl-2 and Bcl-xL proteins respectively (*P*=0.003; *P*=0.043), thus reducing the resistance to apoptosis provided by the antiapoptotic factors of this family (figure 4). In agreement with this observation, we detected an increased in the late apoptotic levels assessed by annexin V and propidium iodide staining in E/R KO clones as compared with control clones (8.99 ± 2.08 vs 2.135 ± 0.065) (*P*<0.05) (Figure 3D). Treatment with Vincristine (1 µM), a drug whose efficacy has been demonstrated in the treatment of ALL by induction of apoptosis in mitotic cells,^58^ induced a greater late apoptotic rate in E/R KO clones as compared with control clones (75.8 ± 9.59 vs 32.7 ± 2.1) (*P*<0.05) (Figure 3E).

**Figure 4.**
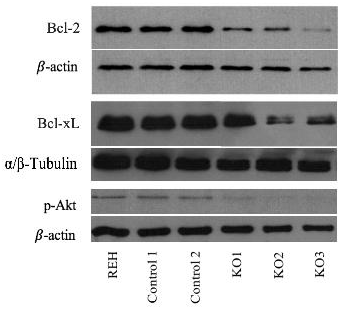
Western blot analysis of E/R targets expression. Lower phospho-Akt (60 kDa), BCL-XL (30 kDa) and BCL-2 (28 kDa) expression levels were observed in all E/R KO clones compared with parental cell line (REH) and control clones.

### Abrogation of ETV6/RUNX1 expression enhances sensitivity to the PI3K inhibitor Copanlisib

Deregulation of *RGS16* or *PTPKR* genes also suggested the alteration of the PI3K/AKT/mTOR pathway. Several studies have already suggested that E/R may be key in the maintenance of the leukaemic phenotype through the activation of different pathways, including the PI3K/AKT/mTOR pathway,^13,59^ resulting in proliferation and cell survival of leukaemic cells. Akt phosphorylation levels measured through WB showed a reduction of 90% in Akt activity in the KO clones relative to REH cells and control clones (*P*=0.003), suggesting the decrease in PI3K/Akt/mTOR activity as a result of the elimination of the expression of E/R (Figure 4).

A large proportion of relapsing positive E/R patients become resistant to GCs such as Prednisolone, widely used in ALL treatment.^8^ Fuka’s group demonstrated that the use of PI3K inhibitors can sensitize positive E/R cells to GCs.^13^

After verifying a lower activation levels of PI3K/Akt/mTOR pathway with the elimination of E/R expression, we aimed to test if these cells responded in the same way to PI3K inhibitors. For that, we used Copanlisib, a PI3K inhibitor with inhibitory activity predominantly against the PI3K-alpha and PI3K-delta isoforms.^60,61^ Treatment with Copanlisib (10 mM) resulted in higher decrease of viability in E/R KO clones compared with REH cells and controls clones (Figure 5A). To verify if Copanlisib was actually inhibiting the PI3K/Akt/mTOR pathway, we measured the phosphorylation levels of Akt and mTOR by WB, before and after treatment. We observed that the phosphorylation levels of both proteins decreased after treatment (Figure 5B).

**Figure 5.**
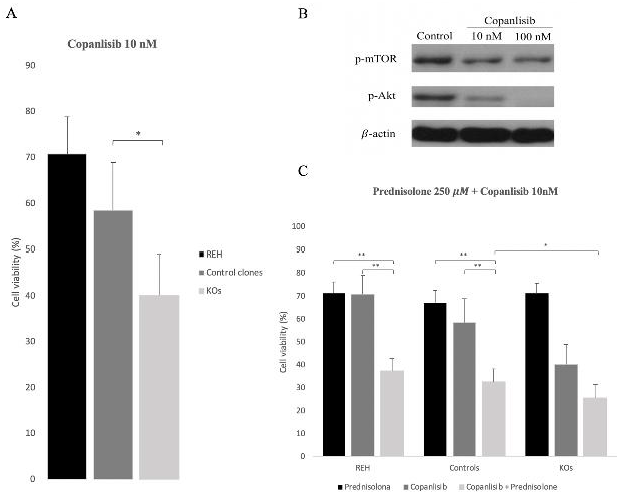
Cell viability and protein expression measured after Copanlisib / Prednisolone treatment. (A) Cell viability was measured by the MTT proliferation assay after treatment (192 h) with Copanlisib (10 nM). E/R KO clones (light grey square line) showed a higher sensitivity to Copanlisib than REH cells (black square line) and control clones (grey square line). This graph represents the average of three independent experiments and in turn the average of the 2 control clones and the 3 KO clones. (B) p-Akt (60 kDa) and p-mTOR (289 kDa) expression levels decreased after treatment with Copanlisib. (C) Prednisolone (black square line), Copanlisib (dark grey square line) and Copanlisib plus Prednisolone combination (grey square line) were tested in the different clones. The relative cell viability was calculated as the percentage of untreated cells. \**P≤*0.05; \*\**P*≤0.005 (unpaired *t*-test).

On the other hand, treatment with Prednisolone (250 µM) was comparable to the effect of Copanlisib on E/R positive cells. We did not observe a higher decrease of cell viability in E/R KO clones as compared with REH cells and control clones (Figure 5C). A joint exposure of Copanlisib (10 nM) and Prednisolone (250 µM) showed a decrease of cell viability in REH and control clones expressing *E/R* fusion gene as compared with Prednisolone and Copanlisib alone. Furthermore, we also observed greater reduction of cell viability in E/R KO clones as compared with REH cells and control clones (Figure 5C).

### E/R repression impairs the tumourigenic capacity *in vivo*

In order to determine the effects of E/R expression abrogation *in vivo*, 16 NSG mice were subcutaneously injected with REH cells or control clone (left flank) and KO clones (right flank). Only 6 mice injected with KO clones developed tumour growth on the right flank (6/16), whereas all those injected with REH or control clone developed a tumour (16/16). In the first group (REH cells vs KO2 clone), none of flanks injected with KO2 developed tumour (mean mass: 0 mg ± 0 vs 4872.5 mg ± 1323; *P*=0.029). In the second group (REH cells vs KO3 clone), only one of flanks injected with KO3 developed a tumour (1/4). This tumour was significantly smaller than those generated from REH cells (mean mass: 40 mg ± 69.3 vs 4212.5 mg ± 1663.9; *P*=0.029). In the third group (control clone vs KO1), we observed tumour growth in 2/4 flanks of mice injected with KO1. These tumours were significantly smaller than those generated from control clone (mean mass: 483 mg ± 354.4 vs 2470 mg ± 872.5 vs *P*=0.041). In the same way, 3/4 mice develop tumours from KO2 in the group 4 (control clone vs KO2), but these tumours were significantly smaller than those generated from control clone (mean mass: 355 mg ± 293.6 vs 2255 mg ± 1215.6; *P*=0.029) (Figure 6A and Supplementary Figure 4). In general, subcutaneous tumours generated from E/R KO cells were significantly smaller than those produced by REH cells or control clone (mean mass: 202 mg ± 298.9 vs 4542.5 mg ± 1539; *P*<0.001// vs 2347.1 mg ± 1087.2; *P*≤0.001) (Figure 6B).

**Figure 6.**
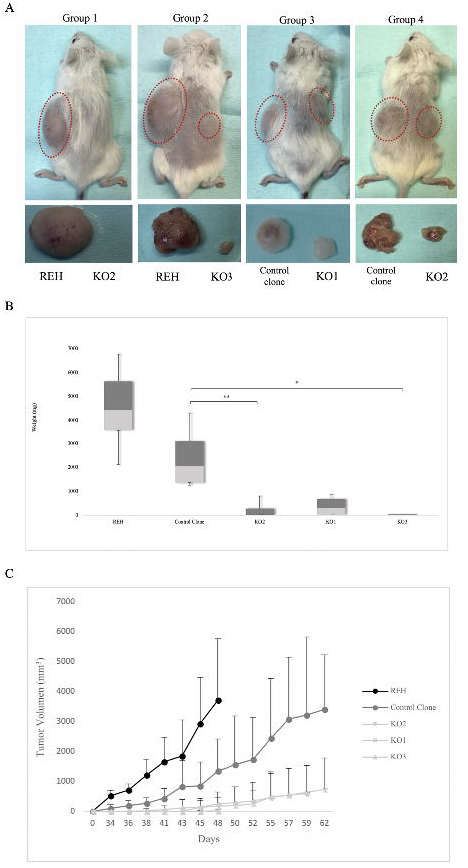
In vivo effects of CRISPR-mediated editing of the *E/R* oncogene. (A) External appearance of mice and developed tumours 48 - 62 days after subcutaneous cell injection. Tumours formed by KO clones (right flank) were smaller than those induced by REH cells or control clones (left flank). (B) Evolution of tumour growth measured every 2-3 days until the moment in which mice were sacrificed. (C) Representation of the mean tumour size corresponding to each clone, independently of the group. \**P≤*0.05; \*\**P≤*0.005 (unpaired *t*-test).

In addition, significant differences were observed in the time of appearance of the tumours. Those tumours generated through the KO clones appeared around day 42 (mean: 46.5 ± 7.8), unlike those generated by the REH cells or the control clone, which appeared around day 29 (mean: 29 ± 4.2; *P*=0.001) and day 36 (mean: 36.6 ± 5.34; *P*=0.03) respectively (Figure 6C).

Histopathological analysis of representative tumours from each group of mice revealed a higher number of mitotic figures in tumours from REH (52 vs 20, *P*=0.017) and control clones (62 vs 20; *P*=0.006) as compared to tumours from KOs clones. In KOs tumours, but not in REH and control clone tumours, we also observed the "starry sky" (macrophages containing dead apoptotic tumour cells) (Supplementary figure 5). No other morphological changes between tumours were observed.

## DISCUSSION

In this study, we generated an E/R KO model in an ALL cell line carrying the t(12;21) in order to assess how the loss of fusion expression affects the tumour cells. This translocation is the most common in children with ALL^1,2^ and encodes a chimeric transcription factor that converts *RUNX1* from a transcriptional modulator to a transcriptional repressor of *RUNX1* target genes.^62^ The expression of E/R results in the generation of a persistent pre-leukaemic clone, which requires of secondary genetic abnormalities to converts to ALL.^57,63^ However, the implication of the E/R expression in the maintenance of the disease is not quite clear. Previous studies have already silenced the fusion protein by using RNAi and shRNA, but they did not completely eliminate the levels of the protein. In addition, these studies showed contradictory results, which suggest that more studies are needed to help elucidate the true oncogenic potential of the fusion protein.^12,13^

The recent evolution of genetic editing techniques with the CRISPR/Cas9 system has allowed, among others, the generation of KO gene models, helping us to better understanding the biology of diseases such as ALL.^64^ We used CRISPR/Cas9 system to completely eliminate the expression of the E/R fusion protein. The loss of fusion expression was checked, observing an absence of mRNA coding for E/R in the KO2 and KO3 clones and a leaky expression in KO1 clone. We hypothesize that this loss of mRNA was due to nonsense-mediated mRNA decay mechanism.^65^ In this way we generated a powerful model on which to study the effect of the elimination of the fusion gene on leukaemic cells. These cells maintain a series of secondary alterations that triggered the leukaemia, similar as occurs in patients.

Transcriptome analysis of different clones showed a huge number of genes significantly deregulated after *E/R* abrogation. A greater number of upregulated genes was observed which agrees with the repressive activity of the *E/R* fusion gene.^62,66^ The most of genes significantly deregulated are involved in different cellular processes such us differentiation and lymphoid activation, apoptosis, cell signaling and cell migration. These results are in agreement with previous studies, in which was demonstrated the implication of E/R in the cellular processes that may be maintaining the leukaemic state of the tumour cells.^13,57,59,66^ The expression of some of these genes has been described as a specific signature of E/R positive ALL patients.^32,33^

Within of the top50 of the significantly deregulated genes in our study we observed a series of downregulated genes after *E/R* abrogation such as *CXCR7*, *LCK* and *PTPGR* and other upregulated genes such as *Mir-146*, *RXRA*, *RGS16*, *TP63* and *IL7R*. The reverse deregulation of these genes, often seen in ALL patients, leads to increased migratory, tumour activation or chemotherapy resistance effects.

Very subtle changes were observed in cell proliferation and cell cycle distribution after *E/R* abrogation *in vitro*, in agreement with previous studies.^12,13^ However, E/R KO clones showed a significantly lower proliferation rate when they were co-cultured with MSC. Mesenchymal cells have been shown to play a key role in the development and evolution of ALL^67,68^ and Bonilla et al. (2019) observed in a recently study that MSCs induce greater cell adhesion, higher proliferation ratio and greater migration capacity to REH cells.^69^ Our data show that the *E/R* fusion gene therefore participates in the interaction of leukaemic cells with the microenvironment and the loss of *E/R* fusion gene expression reverses the proliferative capacity that MSCs confer on leukaemia cells. E/R KO clones also showed a higher late apoptosis rate, demonstrating that the fusion gene regulates the expression of antiapoptotic factors that protect leukaemic cells from apoptosis. Death levels were also higher in E/R KO clones after treatment with Vincristine.

On the other hand, we wanted to check if the non-expression of E/R and consequently the loss of activation of the PI3K/Akt/mTOR pathway, was able to sensitize the cells to PI3K inhibitors. The use of PI3K inhibitors alone has shown to be an effective treatment in E/R positive cells. Furthermore, the activity of these inhibitors in combination with Prednisolone, a GC widely used in the treatment of ALL, has been shown to decrease the resistance offered by positive E/R cells to GCs. In our study, we observed that use of Copanlisib, a PI3K inhibitor, achieved a significantly decrease of cell viability in E/R KO clones as compared with E/R positive cells. We also observed that treatment with Copanlisib achieved the sensitization to Prednisolone in E/R positive cells as Fuka’s study described.^13^ However, in our study, this sensitization was even greater in E/R KO cells. Therefore, our data demonstrate that the fusion gene may be a good therapeutic target with which to improve the drug sensitivity of positive E/R cells.

Finally, we wanted to check if E/R abrogation also decreased the tumour potential of cells *in vivo*. For that, a xenograft model was generated by injecting these cells into immunosuppressed mice, taking the injection of REH cells or a control clone on the opposite flank as control. Mice injected with KO clone cells did not generated tumours or generated smaller tumours than those generated by REH cells or control clone. The higher rate of mitotic activity in REH and control tumours observed through the histopathology analysis explains the greater growth of these tumours and reveals a greater tumoural capacity of these cells carrying *E/R* fusion gen *in vivo*.

Together these data suggest that the *E/R* fusion gene has a key role in the maintenance of the leukaemic phenotype. By eliminating E/R expression, the cells lost tumourigenic capacity, becoming more sensitive to drugs and reducing their oncogenic capacity *in vitro* and *in vivo*. Therefore, although more studies are needed to elucidate the mechanism of action of the fusion gene, this study demonstrates that it could be a possible therapeutic target to design new drugs that prevent the correct expression of this protein.

## AUTHORS CONTRIBUITION

AM, IGT, RB and JMHR designed research. AM performed the experiments, compiled data and drafted the manuscript. JLO was responsible for mouse experiments and drug tests. JHS was responsible for sequencing experiments. VAP carried out additional experimental work. IGT, RB, JMHR, JLO, VAP and TG helped interpret the results and write the manuscript. AM wrote the paper with input from all authors.

## ACKNOWLEDGEMENTS

We thank Sara González, Irene Rodríguez, Maribel Forero-Castro, Ana Marín-Quílez, María Herrero-García, Almudena Martín-Martín, Sandra Santos, Cristina Miguel, from the Cancer Research Center of Salamanca, Spain. We are grateful to Ángel Prieto and Ana I García, María Luz Sánchez and María Carmen Macías from the Microscopy Unit, Cytometry Unit and Molecular Pathology Unit, respectively, from the Cancer Research Center of Salamanca for the technical assistance. We thank Luis Muñoz and all the members from the Animal Experimentation Research Center from the University of Salamanca. We also thank Mercedes Garayoa for providing us with the HS-5 cell line.

## FUNDING

This work was financially supported in part by a grant from the Consejería de Educación, Junta de Castilla y León, Fondos FEDER (SA085U16, SA271P18), and the Regional Council of Castilla y León SACYL, (GRS 1847/A/18), Fundación Castellano Leonesa de Hematología y Hemoterapia (FUCALHH 2017), Proyectos de investigación en Biomedicina, gestión sanitaria y atención sociosanitaria del IBSAL (IBY17/00006), Fundación Memoria Don Samuel Solórzano Barruso, Centro de Investigación Biomédica en Red de Cáncer (CIBERONC CB16/12/00233), a grant to JLO from the University of Salamanca (“Contrato postdoctoral programa II 2017-18”), and a grant to AM from the Junta Provincial de Salamanca of the Asociación Española Contra el Cáncer (AECC).

## COMPETING INTERESTS

The authors declare that they have no competing interests

