## Supplementary for "*Etv6/Runx1* Fusion Gene Abrogation Decreases The Oncogenic Potencial Of Tumour Cells In A Preclinical Model Of Acute Lymphoblastic Leukaemia"

*These authors share senior authorship.

**Corresponding author:**

Jesús-María Hernández-Rivas

**Running title:** *E/R* maintains the oncogenic potential of ALL cells

**Keywords**

Acute lymphoblastic leukemia, ETV6/RUNX1, CRISPR/Cas9, Genome edition.

**Supplementary tables and figures**

**Supplementary table 1. sgRNA designed against *E/R* fusion sequence.** Two custom-designed single guide RNAs (sgRNAs) were designed to genetically inactivate the *E/R* oncogene. These specific sgRNAs direct Cas9 to the *E/R* fusion sequence. G1 sgRNA, directed at the end of exon 5 of *ETV6* and G2 directed to intronic region before the fusion point between *ETV6* and *RUNX1*.

|  | Forward | Reverse |
| --- | --- | --- |
| G 1 | caccgGCCTAATTGGGAATGGTGCG | aaacCGCACCATTCCCAATTAGGCc |
| G 2 | caccgAAGAGCACGCCATGCCCATT | aaacAATGGGCATGGCGTGCTCTTc |

**Supplementary table 2. Possible off-targets of the sgRNAs.** The possible off-targets of the sgRNAs used obtained from “Breaking Cas” website (<http://bioinfogp.cnb.csic.es/tools/breakingcas/>) were checked by PCR and sanger sequencing. Two pairs of oligos were designed for each off-target region.

| Gen | Score | Forward | | Reverse |
| --- | --- | --- | --- | --- |
| PTPN21 | 0 | AGTGTAATTTGGGAAACAGCCCT | TTGGGGCTTTTCCCACCTCAA | |
| BCL9L | 0.3 | AGAGAATGGATCTGGGAGGGA | AGGCGAGGCAGTTGCAGTGTA | |
| BEND4 | 0.1 | GGCATCAGGTAGCCAACGTTC | CAGGATCAATGGATTTGTCACAG | |
| VSIR | 0 | AAAGGGTCCAGAGAAGAGAGG | CTGGGGACGGAGCAAAACTTT | |

**Supplementary table 3. Genes significantly deregulated after *E/R* fusion gene abrogation.** List of 342 genes significantly deregulated after *E/R* fusion gene abrogation sorted according to the decreasing value of the FC. The TP50 of deregulated genes are shaded in grey. In the following columns from left to right, the mean of normalized counts of all samples (baseMean); log2-Fold Change (log2FC); log2FC unshrunk, P-value and adjusted P value.

| Gene | baseMean | log2FC | log2FCunshrunk | Pvalue | Padj |
| --- | --- | --- | --- | --- | --- |
| *DRD5* | 154.4 | -2.34959 | -3.92883 | 0 | 0 |
| *ATP10A* | 231.1 | -2.07428 | -3.35631 | 0 | 0 |
| *SGIP1* | 70.3 | -2.0645 | -3.86631 | 0 | 0 |
| *BIRC3* | 80.1 | 1.98755 | 4.09318 | 0 | 0 |
| *ACKR3* | 66.8 | -1.98193 | -3.84476 | 0 | 0 |
| *EMR2* | 80.4 | 1.97712 | 3.98976 | 0 | 0 |
| *FAM189A1* | 55.5 | -1.80486 | -4.18028 | 0 | 1.97E-13 |
| *MIR146A* | 278.6 | 1.77216 | 2.54174 | 0 | 1E-15 |
| *OPTN* | 315.4 | 1.7413 | 2.24832 | 0 | 0 |
| *LHX6* | 127.2 | 1.72519 | 3.8055 | 4E-15 | 3.17E-12 |
| *NBPF3* | 35.4 | -1.67532 | -4.9272 | 1.6E-14 | 1.23E-11 |
| *GCSAM* | 1072 | -1.67221 | -1.90226 | 0 | 0 |
| *PGPEP1* | 94.7 | -1.66199 | -3.62221 | 3.7E-14 | 2.72E-11 |
| *KCNA2* | 143.1 | -1.65663 | -2.17481 | 0 | 1E-15 |
| *RP11-622K12.1* | 367.2 | 1.59169 | 2.28594 | 2E-15 | 1.94E-12 |
| *RASD2* | 209.2 | -1.47352 | -2.0491 | 6.1E-14 | 4.04E-11 |
| *CDH23* | 265.2 | -1.44672 | -1.96584 | 6E-14 | 4.04E-11 |
| *ALOX5* | 89.6 | 1.37691 | 2.54469 | 2.65E-10 | 1.38E-07 |
| *SORBS2* | 882 | -1.364 | -1.63045 | 0 | 4.4E-14 |
| *HYDIN* | 83.6 | 1.36022 | 3.17093 | 5.3E-10 | 2.67E-07 |
| *GPR17* | 119.4 | -1.33855 | -2.68649 | 1.05E-09 | 4.79E-07 |
| *MOXD1* | 162.7 | 1.33314 | 3.05082 | 1.17E-09 | 5.15E-07 |
| *ARX* | 64.4 | -1.30499 | -2.44801 | 2.28E-09 | 9.76E-07 |
| *KCNE3* | 249.5 | -1.30368 | -1.95828 | 2.64E-10 | 1.38E-07 |
| *HOXA13* | 234.3 | 1.29849 | 1.52883 | 0 | 1.13E-13 |
| *LCK* | 214.4 | -1.29552 | -2.04607 | 7.9E-10 | 3.84E-07 |
| *RELB* | 271.2 | 1.28523 | 1.51994 | 0 | 4.11E-13 |
| *DDR1* | 207.4 | 1.23577 | 1.58447 | 9.25E-12 | 5.62E-09 |
| *SCN3A* | 361.8 | 1.23448 | 1.91793 | 3.86E-09 | 1.52E-06 |
| *FRMD4B* | 807.8 | -1.22731 | -1.62866 | 8.07E-11 | 4.52E-08 |
| *LIMS2* | 495.1 | -1.20713 | -2.10548 | 2.5E-08 | 7.02E-06 |
| *RGS1* | 221.1 | -1.19302 | -1.85261 | 1.24E-08 | 4.03E-06 |
| *CALML6* | 75.1 | 1.19083 | 1.84701 | 1.21E-08 | 4E-06 |
| *FZD3* | 83 | 1.179 | 2.86812 | 6.42E-08 | 1.49E-05 |
| *ARHGAP42P2* | 187.8 | -1.17718 | -2.04583 | 5.34E-08 | 1.25E-05 |
| *RP11-251M1.1* | 558.7 | 1.1701 | 1.89706 | 3.88E-08 | 9.91E-06 |
| *RXRA* | 695.7 | 1.16993 | 1.68558 | 6.77E-09 | 2.47E-06 |
| *RGS16* | 431.5 | 1.16993 | 4.27179 | 1.85E-08 | 5.52E-06 |
| *HOTTIP* | 40.8 | 1.16894 | 2.4038 | 1.01E-07 | 2.26E-05 |
| *FBN3* | 196.1 | -1.16298 | -1.45832 | 3.86E-11 | 2.25E-08 |
| *RP11-16L9.4* | 439.2 | 1.15288 | 1.66285 | 1.14E-08 | 3.86E-06 |
| *HLX* | 101.3 | 1.14724 | 1.59611 | 5.25E-09 | 1.97E-06 |
| *TP63* | 318.4 | 1.1262 | 1.88183 | 1.55E-07 | 3.14E-05 |
| *TLR7* | 61.4 | 1.12406 | 2.8608 | 2.25E-07 | 4.37E-05 |
| *PTPRG-AS1* | 73.4 | -1.11862 | -3.58436 | 1.17E-07 | 2.44E-05 |
| *FAM78A* | 548.8 | 1.11483 | 1.43424 | 9.77E-10 | 4.59E-07 |
| *BCAS1* | 210.8 | -1.11072 | -1.8555 | 2.28E-07 | 4.38E-05 |
| *PLS1* | 62.8 | 1.10389 | 2.20117 | 4.87E-07 | 8.16E-05 |
| *AQP5* | 229.5 | 1.09449 | 1.83256 | 3.47E-07 | 6.1E-05 |
| *PDE10A* | 95.9 | -1.08807 | -2.11786 | 6.94E-07 | 0.000109 |
| *CSF1* | 29.6 | 1.07667 | 3.30357 | 4.18E-07 | 7.16E-05 |
| *TTC28* | 422.8 | -1.06872 | -1.42756 | 1.9E-08 | 5.53E-06 |
| *EPHB6* | 332.1 | 1.05097 | 1.40655 | 3.42E-08 | 8.9E-06 |
| *GIMAP6* | 104.7 | -1.04917 | -2.08309 | 1.74E-06 | 0.000237 |
| *CNR2* | 151.2 | -1.04836 | -1.46725 | 1.15E-07 | 2.43E-05 |
| *SGK1* | 203.2 | 1.04539 | 1.4315 | 7.51E-08 | 1.71E-05 |
| *TRPM2* | 307.5 | 1.04389 | 1.56381 | 4.22E-07 | 7.16E-05 |
| *SLC25A4* | 375.9 | -1.04233 | -1.35587 | 1.71E-08 | 5.4E-06 |
| *SERPINB1* | 449.6 | -1.03395 | -1.48499 | 2.89E-07 | 5.4E-05 |
| *ADCY7* | 789.1 | 1.02767 | 1.67585 | 1.47E-06 | 0.000206 |
| *EPN2* | 211.7 | -1.02564 | -2.01505 | 2.94E-06 | 0.000382 |
| *NPL* | 61.1 | 1.01372 | 1.61799 | 1.64E-06 | 0.000226 |
| *RP11-1055B8.4* | 129.1 | 1.00869 | 1.41433 | 3.47E-07 | 6.1E-05 |
| *CD52* | 5748.7 | -1.00638 | -1.07155 | 0 | 0 |
| *C10orf10* | 193.8 | -1.00598 | -1.707 | 3.06E-06 | 0.000394 |
| *MYO7B* | 130 | -1.00505 | -2.57867 | 3.37E-06 | 0.000419 |
| *N4BP3* | 1631.4 | 1.0044 | 1.34839 | 1.49E-07 | 3.05E-05 |
| *TRNP1* | 843.7 | -1.00257 | -1.36303 | 2.17E-07 | 4.27E-05 |
| *RP11-443A13.5* | 571.4 | 1.00152 | 1.21962 | 2.54E-09 | 1.06E-06 |
| *LRP4* | 64.7 | 0.99177 | 2.46045 | 4.97E-06 | 0.000603 |
| *TSPAN9* | 526.5 | -0.99105 | -1.11836 | 1.14E-12 | 7.23E-10 |
| *TMEM64* | 260.3 | 0.98891 | 1.45127 | 1.27E-06 | 0.000182 |
| *TCF7* | 151 | -0.98381 | -1.32807 | 2.96E-07 | 5.46E-05 |
| *MMP14* | 192.5 | 0.98032 | 1.80139 | 7.14E-06 | 0.000846 |
| *FAM107B* | 10387.3 | -0.97505 | -1.19805 | 1.14E-08 | 3.86E-06 |
| *PLEK* | 1078.2 | 0.96863 | 1.28742 | 3.1E-07 | 5.65E-05 |
| *NOL4* | 214.5 | 0.96397 | 1.17625 | 1.07E-08 | 3.8E-06 |
| *CAV1* | 297 | -0.95749 | -1.27142 | 4.08E-07 | 7.08E-05 |
| *AGR2* | 38.8 | 0.95646 | 5.4769 | 9.71E-07 | 0.000149 |
| *RGMA* | 1017.8 | -0.95634 | -1.18978 | 4.04E-08 | 1.02E-05 |
| *RASSF4* | 289 | -0.95437 | -1.75826 | 1.25E-05 | 0.001381 |
| *LCN6* | 249.8 | 0.95321 | 1.81714 | 1.36E-05 | 0.001474 |
| *RP11-1055B8.7* | 6983 | 0.94416 | 1.00376 | 0 | 0 |
| *PTMS* | 72.4 | 0.94242 | 1.9579 | 1.77E-05 | 0.001864 |
| *CYTH4* | 524.2 | 0.9396 | 1.24137 | 5.87E-07 | 9.41E-05 |
| *KLHDC7B* | 22.2 | 0.93929 | 2.83172 | 9.63E-06 | 0.001105 |
| *HAP1* | 3579.1 | -0.936 | -1.13741 | 2.31E-08 | 6.59E-06 |
| *RHOH* | 1146.9 | -0.9267 | -1.23334 | 1.02E-06 | 0.000153 |
| *IRF8* | 803.1 | -0.92338 | -1.25995 | 1.95E-06 | 0.000263 |
| *LARGE* | 415.8 | -0.91806 | -1.3959 | 1.02E-05 | 0.001156 |
| *LAMB4* | 79.9 | -0.9156 | -1.67425 | 2.72E-05 | 0.002662 |
| *AK7* | 1333.1 | -0.91269 | -1.0905 | 1.8E-08 | 5.52E-06 |
| *TBXAS1* | 369.3 | 0.9111 | 1.09726 | 3.12E-08 | 8.26E-06 |
| *PLEKHB1* | 67.2 | 0.90934 | 1.81052 | 3.42E-05 | 0.00322 |
| *SPTBN4* | 89.6 | -0.90781 | -1.24563 | 3.11E-06 | 0.000397 |
| *CXXC5* | 1769.6 | 0.90625 | 1.05615 | 3.28E-09 | 1.33E-06 |
| *GUCY1A3* | 222.9 | -0.89857 | -1.50261 | 2.9E-05 | 0.00282 |
| *GPR142* | 27.8 | -0.89415 | -2.85582 | 1.96E-05 | 0.001998 |
| *ZNF264* | 134.7 | 0.89389 | 1.26566 | 7.45E-06 | 0.000869 |
| *SNTA1* | 325.1 | -0.88594 | -1.09918 | 3.25E-07 | 5.86E-05 |
| *SV2A* | 972.4 | 0.88563 | 1.08285 | 1.7E-07 | 3.39E-05 |
| *ZNF469* | 158.6 | 0.88264 | 1.58403 | 5.07E-05 | 0.004447 |
| *PIGM* | 310.8 | 0.88148 | 1.10151 | 4.97E-07 | 8.23E-05 |
| *AC090044.2* | 95.8 | 0.87855 | 1.7981 | 6.3E-05 | 0.005364 |
| *SEMA7A* | 122.3 | 0.87808 | 1.47324 | 4.44E-05 | 0.004041 |
| *NTN3* | 154.2 | 0.87613 | 1.09977 | 6.78E-07 | 0.000107 |
| *LINC01013* | 150 | -0.87602 | -1.36521 | 3.06E-05 | 0.002955 |
| *CROCC* | 298.9 | 0.87521 | 1.03825 | 4.13E-08 | 1.02E-05 |
| *MMP15* | 449.3 | -0.8698 | -1.13171 | 2.58E-06 | 0.000344 |
| *RP11-1055B8.6* | 283 | 0.86919 | 1.226 | 1.27E-05 | 0.001392 |
| *EMP2* | 378.1 | 0.86759 | 1.09257 | 9.99E-07 | 0.000152 |
| *RP11-431N15.2* | 20.1 | 0.86663 | 5.26703 | 6.26E-06 | 0.000747 |
| *ZNF528* | 133.8 | 0.8636 | 1.48223 | 6.46E-05 | 0.005392 |
| *FCGR2A* | 66.4 | 0.86013 | 1.72152 | 8.92E-05 | 0.006923 |
| *NEIL1* | 67.7 | 0.85161 | 1.49228 | 8.63E-05 | 0.006797 |
| *SIT1* | 183.8 | 0.85063 | 1.50487 | 9.1E-05 | 0.006981 |
| *NT5E* | 323.3 | -0.85004 | -1.0867 | 2.66E-06 | 0.000352 |
| *CECR2* | 268.9 | 0.84865 | 1.17241 | 1.5E-05 | 0.001603 |
| *SERPINB6* | 560.5 | -0.84378 | -1.0843 | 3.67E-06 | 0.000453 |
| *SIDT1* | 263.9 | -0.84107 | -1.39962 | 8.96E-05 | 0.006923 |
| *AFF2* | 979.3 | 0.83909 | 1.10828 | 8.18E-06 | 0.000947 |
| *TEAD3* | 73 | 0.83854 | 1.4096 | 9.67E-05 | 0.007272 |
| *CD302* | 207.1 | 0.83694 | 1.39459 | 9.73E-05 | 0.007272 |
| *C1QTNF4* | 454.5 | -0.83462 | -1.02567 | 1.03E-06 | 0.000153 |
| *PTPN7* | 496.1 | -0.83337 | -1.42407 | 0.000115 | 0.008254 |
| *LBH* | 1252.6 | -0.8326 | -1.02589 | 1.22E-06 | 0.000178 |
| *RIC3* | 24 | 0.83244 | 4.0662 | 1.94E-05 | 0.001998 |
| *SEMA4A* | 185.8 | 0.83085 | 1.17682 | 3.18E-05 | 0.003026 |
| *NRBP2* | 246.9 | 0.82947 | 1.18658 | 3.71E-05 | 0.00344 |
| *TSKU* | 93.4 | 0.82585 | 1.35517 | 0.000112 | 0.008122 |
| *CD40* | 168.6 | 0.82537 | 1.01695 | 1.47E-06 | 0.000206 |
| *DBNDD1* | 403.2 | -0.82471 | -1.16418 | 3.51E-05 | 0.00328 |
| *KCNJ4* | 36.7 | -0.82267 | -1.86593 | 0.000169 | 0.011166 |
| *SORBS1* | 27.8 | 0.81941 | 2.04647 | 0.000157 | 0.010539 |
| *IL4I1* | 31.7 | 0.81657 | 2.62006 | 8.98E-05 | 0.006923 |
| *RP11-469H8.6* | 67.9 | 0.81609 | 2.12138 | 0.000151 | 0.010331 |
| *GNG11* | 1694.7 | -0.81545 | -1.06571 | 1.16E-05 | 0.001309 |
| *CAP2* | 128.4 | -0.81443 | -1.23333 | 8.73E-05 | 0.006837 |
| *PLEC* | 410.6 | 0.80672 | 1.04502 | 1.17E-05 | 0.001309 |
| *PDZD7* | 100.6 | 0.80051 | 1.19706 | 0.000104 | 0.007603 |
| *TMEM52* | 235.2 | 0.79981 | 1.61531 | 0.00027 | 0.016797 |
| *AC090044.1* | 372.3 | 0.79949 | 1.33198 | 0.000198 | 0.01259 |
| *ATP13A2* | 773.7 | 0.79809 | 0.98318 | 3.28E-06 | 0.000412 |
| *NPY* | 354.1 | 0.79767 | 1.1838 | 0.000104 | 0.007611 |
| *SLC51A* | 923.8 | -0.79604 | -1.11872 | 6.19E-05 | 0.005302 |
| *PSD2* | 32.5 | 0.78983 | 1.91702 | 0.000279 | 0.017214 |
| *CAST* | 535.9 | 0.78854 | 1.09765 | 6.42E-05 | 0.005392 |
| *PRR5L* | 357.9 | -0.785 | -0.9029 | 1.1E-07 | 2.39E-05 |
| *ARHGEF12* | 926.6 | -0.78172 | -0.87228 | 5.27E-09 | 1.97E-06 |
| *C20orf197* | 21.9 | 0.78168 | 2.91678 | 0.000115 | 0.008254 |
| *ZNF704* | 1650.9 | -0.78124 | -0.89744 | 1.15E-07 | 2.43E-05 |
| *DAB2IP* | 529.9 | 0.77986 | 0.99995 | 1.79E-05 | 0.001872 |
| *TMEM173* | 173.5 | 0.77921 | 1.1944 | 0.00019 | 0.012234 |
| *IGHD* | 430.3 | 0.77753 | 1.17758 | 0.000182 | 0.011865 |
| *ACTN1* | 1971.1 | 0.7764 | 0.88027 | 4.2E-08 | 1.02E-05 |
| *RP3-455J7.4* | 112.6 | 0.7684 | 1.51577 | 0.000463 | 0.025573 |
| *TMEM229B* | 179.1 | 0.76807 | 1.38713 | 0.000428 | 0.024342 |
| *NAT8L* | 59 | 0.7666 | 1.47598 | 0.000471 | 0.02581 |
| *RBMS2* | 105.9 | 0.76601 | 1.02433 | 5.69E-05 | 0.004937 |
| *SPNS2* | 96.1 | -0.76441 | -1.13302 | 0.000197 | 0.01259 |
| *FAM65B* | 1931.9 | -0.76332 | -1.03087 | 7.24E-05 | 0.005959 |
| *KCTD12* | 44.2 | 0.7633 | 1.79551 | 0.00046 | 0.025497 |
| *LOXHD1* | 82 | -0.76287 | -2.05591 | 0.000363 | 0.021387 |
| *ARL4C* | 117.7 | -0.7613 | -1.60277 | 0.000521 | 0.027621 |
| *PDLIM1* | 3779.7 | -0.75931 | -1.07166 | 0.00014 | 0.009802 |
| *ZNF169* | 102.5 | 0.75727 | 1.06339 | 0.000135 | 0.009626 |
| *TPM2* | 503.6 | 0.75349 | 0.95242 | 2.39E-05 | 0.002421 |
| *PFN2* | 420.2 | 0.75328 | 1.19692 | 0.000376 | 0.022024 |
| *WFS1* | 101.5 | 0.75075 | 1.18653 | 0.000381 | 0.022203 |
| *LCN10* | 42.8 | 0.74986 | 1.967 | 0.000486 | 0.026329 |
| *LILRB4* | 30.2 | -0.74518 | -2.02167 | 0.000494 | 0.02656 |
| *NCALD* | 46.4 | -0.74335 | -1.48403 | 0.000709 | 0.034244 |
| *ID2* | 152.8 | 0.74327 | 1.53331 | 0.00071 | 0.034244 |
| *CAMK4* | 217.7 | 0.74297 | 0.96783 | 6.03E-05 | 0.0052 |
| *PDGFA* | 327.7 | 0.74222 | 1.18772 | 0.000473 | 0.025829 |
| *TEX14* | 53.7 | 0.74155 | 2.36737 | 0.000355 | 0.02103 |
| *SPRY1* | 230.1 | -0.73794 | -0.92144 | 2.53E-05 | 0.002512 |
| *STIM1* | 473.7 | -0.73641 | -0.97332 | 9.19E-05 | 0.00701 |
| *SPARC* | 101.2 | 0.7362 | 2.12186 | 0.000497 | 0.026604 |
| *CACNA2D1* | 75.1 | 0.73588 | 1.23955 | 0.000631 | 0.031553 |
| *ITGB2* | 713.9 | 0.73385 | 0.972 | 0.000101 | 0.007414 |
| *CTD-3018O17.3* | 68.5 | 0.73343 | 2.08948 | 0.000537 | 0.028172 |
| *ANO1* | 25.8 | -0.73342 | -1.94635 | 0.000632 | 0.031553 |
| *C20orf194* | 216.2 | 0.73325 | 0.93779 | 5.16E-05 | 0.004504 |
| *MS4A4A* | 134.3 | 0.73223 | 0.95342 | 7.59E-05 | 0.006175 |
| *GARNL3* | 45.5 | 0.73071 | 1.25472 | 0.000723 | 0.034641 |
| *CARD9* | 84.4 | 0.72843 | 1.27716 | 0.000792 | 0.037227 |
| *LINC00114* | 50.9 | 0.72708 | 1.56987 | 0.000912 | 0.041384 |
| *GALNT10* | 153.9 | 0.72678 | 1.17004 | 0.000634 | 0.031553 |
| *C1orf222* | 60.1 | 0.7267 | 3.38097 | 0.00016 | 0.010692 |
| *RASGRP2* | 5671 | 0.72471 | 0.80094 | 1.86E-08 | 5.52E-06 |
| *RP11-830F9.7* | 568.5 | 0.72421 | 0.97573 | 0.000161 | 0.0107 |
| *MAP3K15* | 84 | 0.7234 | 2.14555 | 0.000581 | 0.030165 |
| *TACC2* | 14.2 | -0.7217 | -2.80884 | 0.000312 | 0.018838 |
| *CORO2A* | 263.9 | -0.72125 | -1.07485 | 0.000465 | 0.025585 |
| *TNFAIP8* | 784.8 | -0.72046 | -0.80155 | 5.25E-08 | 1.25E-05 |
| *PKIG* | 180.6 | 0.7152 | 1.42286 | 0.001122 | 0.048384 |
| *TLR3* | 112 | 0.71488 | 1.14196 | 0.000752 | 0.035921 |
| *LMNA* | 28 | 0.71359 | 1.78526 | 0.000974 | 0.043658 |
| *TMPRSS15* | 176.6 | -0.71246 | -1.40711 | 0.001171 | 0.049885 |
| *FAM132B* | 448.9 | -0.71207 | -0.93437 | 0.000138 | 0.009751 |
| *LACC1* | 65.3 | 0.71038 | 1.2899 | 0.00113 | 0.048557 |
| *BRINP2* | 106.2 | -0.70892 | -1.81139 | 0.000999 | 0.044262 |
| *HNRNPA1P37* | 148.6 | -0.70868 | -1.18719 | 0.000988 | 0.04402 |
| *FAIM3* | 190.7 | -0.70773 | -1.18642 | 0.001006 | 0.044414 |
| *ZNF641* | 98.1 | 0.70696 | 1.05068 | 0.000587 | 0.030165 |
| *TTC9* | 223.9 | -0.7064 | -1.07029 | 0.000674 | 0.032979 |
| *ABLIM1* | 225.1 | -0.70592 | -1.07602 | 0.000704 | 0.034244 |
| *C10orf128* | 142.5 | 0.70535 | 1.8123 | 0.00105 | 0.045788 |
| *BTN2A3P* | 99.4 | 0.69936 | 1.01969 | 0.000588 | 0.030165 |
| *PTPN3* | 66.8 | -0.69834 | -1.16774 | 0.001161 | 0.049594 |
| *SERINC5* | 435.8 | -0.69699 | -0.97851 | 0.000449 | 0.02528 |
| *SLC38A1* | 6284.3 | -0.69645 | -0.87412 | 8.05E-05 | 0.006483 |
| *DAPK2* | 666.3 | -0.69437 | -0.83711 | 2.59E-05 | 0.002554 |
| *GIMAP8* | 19.7 | -0.69337 | -2.63213 | 0.000536 | 0.028172 |
| *BMP2K* | 1727.1 | -0.69233 | -0.75947 | 2.78E-08 | 7.64E-06 |
| *HIP1R* | 570 | -0.6917 | -0.84738 | 4.71E-05 | 0.004258 |
| *HTRA3* | 17.2 | 0.6915 | 2.51822 | 0.000634 | 0.031553 |
| *DSTYK* | 220 | 0.6907 | 0.87977 | 0.000126 | 0.009033 |
| *BCL11B* | 316.1 | 0.68927 | 0.99353 | 0.000652 | 0.03221 |
| *PLXNC1* | 284.8 | 0.68588 | 0.78584 | 2.71E-06 | 0.000356 |
| *ORAI2* | 1167.4 | -0.68344 | -0.88109 | 0.000189 | 0.012234 |
| *XBP1* | 5123.5 | -0.68319 | -0.93619 | 0.000456 | 0.025497 |
| *CDH24* | 1294 | 0.68212 | 0.84292 | 7.64E-05 | 0.006185 |
| *CD72* | 1662 | 0.68122 | 0.88181 | 0.000213 | 0.013462 |
| *PCDHGC4* | 364.2 | -0.68111 | -0.82387 | 4.12E-05 | 0.0038 |
| *GREB1* | 1076 | -0.68088 | -0.95199 | 0.000586 | 0.030165 |
| *KCNQ1OT1* | 78.4 | 0.67777 | 1.03449 | 0.001142 | 0.048945 |
| *MBOAT1* | 104.5 | 0.67352 | 1.00355 | 0.001076 | 0.046676 |
| *ZFHX3* | 434.9 | 0.67246 | 0.91572 | 0.00052 | 0.027621 |
| *NEAT1* | 2886.8 | 0.67221 | 0.95417 | 0.000787 | 0.037215 |
| *PCCA* | 700.6 | -0.67141 | -0.84344 | 0.000147 | 0.010087 |
| *LRRC4* | 495.5 | -0.6704 | -0.79996 | 3.38E-05 | 0.003202 |
| *BCL6* | 2112.8 | -0.66649 | -0.72574 | 2.96E-08 | 7.98E-06 |
| *PTPRK* | 5372.9 | -0.66494 | -0.75968 | 4.57E-06 | 0.000559 |
| *ALDH3A2* | 645 | -0.66287 | -0.85065 | 0.000271 | 0.016797 |
| *METRNL* | 199.1 | -0.6628 | -0.81304 | 9.94E-05 | 0.007392 |
| *RP11-715J22.6* | 155.9 | 0.66094 | 0.85659 | 0.000331 | 0.019844 |
| *FMO4* | 18.3 | 0.65854 | 2.85429 | 0.000706 | 0.034244 |
| *MFI2* | 975.1 | 0.65699 | 0.78398 | 4.86E-05 | 0.004367 |
| *CBR3-AS1* | 468.6 | -0.65573 | -0.7299 | 7.64E-07 | 0.000118 |
| *GALNT12* | 180.5 | 0.65362 | 0.79167 | 8.6E-05 | 0.006797 |
| *MECOM* | 542.8 | -0.64174 | -0.83026 | 0.000482 | 0.02623 |
| *GIPC1* | 309.8 | 0.6412 | 0.80317 | 0.000271 | 0.016797 |
| *SIKE1* | 1347.5 | -0.63951 | -0.85494 | 0.00079 | 0.037227 |
| *TAGLN3* | 209.4 | -0.63932 | -0.79078 | 0.000213 | 0.013462 |
| *AC002454.1* | 1316.6 | -0.63931 | -0.76796 | 9.63E-05 | 0.007272 |
| *AATK* | 11.4 | -0.6371 | -3.43338 | 0.000604 | 0.03067 |
| *ZC3H12A* | 275.7 | 0.63261 | 0.82339 | 0.000632 | 0.031553 |
| *MYOM2* | 241.4 | 0.63191 | 0.82335 | 0.000651 | 0.03221 |
| *LAX1* | 250.8 | -0.61615 | -0.79876 | 0.000827 | 0.038338 |
| *RMND5B* | 689.1 | 0.61304 | 0.75966 | 0.000402 | 0.023211 |
| *GPSM1* | 5511.7 | 0.60465 | 0.71777 | 0.000157 | 0.010539 |
| *SSH1* | 1022.2 | -0.60434 | -0.66309 | 1.27E-06 | 0.000182 |
| *DPEP1* | 659.7 | 0.59957 | 3.82265 | 0.000718 | 0.034544 |
| *PECR* | 528.8 | -0.5978 | -0.66384 | 5.37E-06 | 0.000646 |
| *AVEN* | 268.8 | -0.59717 | -0.75211 | 0.000767 | 0.036418 |
| *LIG4* | 1424.3 | -0.59515 | -0.76201 | 0.001043 | 0.045721 |
| *JAM2* | 167.8 | 0.59361 | 0.74641 | 0.000798 | 0.037408 |
| *C12orf23* | 3429.3 | -0.59254 | -0.63665 | 1.07E-07 | 2.36E-05 |
| *HLA-F* | 1579 | 0.59166 | 0.72213 | 0.00046 | 0.025497 |
| *TMEM169* | 243.2 | -0.58601 | -0.73804 | 0.00096 | 0.043177 |
| *PPM1L* | 569 | 0.58504 | 0.69388 | 0.000247 | 0.015533 |
| *GRAMD1B* | 666.5 | -0.58441 | -0.72068 | 0.000674 | 0.032979 |
| *TCL1A* | 14125.8 | -0.58379 | -0.7102 | 0.000508 | 0.027111 |
| *LTBP4* | 1986 | -0.58366 | -0.70366 | 0.000408 | 0.023403 |
| *GABRA3* | 600.3 | 0.58365 | 0.73228 | 0.000946 | 0.042792 |
| *CLSTN3* | 436.5 | 0.58028 | 0.65011 | 1.96E-05 | 0.001998 |
| *OPN3* | 765.2 | -0.57808 | -0.64306 | 1.25E-05 | 0.001381 |
| *INO80C* | 371.9 | 0.57505 | 0.70177 | 0.000659 | 0.03245 |
| *AJUBA* | 747 | 0.57157 | 0.65377 | 8.59E-05 | 0.006797 |
| *LRRC8C* | 2327.4 | -0.57157 | -0.70511 | 0.000891 | 0.040809 |
| *PIK3C3* | 6227 | -0.56954 | -0.63715 | 2.53E-05 | 0.002512 |
| *TSPAN7* | 257.4 | 0.56592 | 0.65737 | 0.000194 | 0.012467 |
| *ANXA2* | 1995 | 0.56427 | 0.62478 | 1.39E-05 | 0.001501 |
| *IL1RAP* | 795.7 | -0.56395 | -0.66952 | 0.000424 | 0.024203 |
| *NFKB2* | 609.9 | 0.56343 | 0.67862 | 0.000628 | 0.031553 |
| *BLVRB* | 781.5 | -0.56098 | -0.62694 | 3.12E-05 | 0.002991 |
| *IER2* | 3387.6 | 0.56098 | 0.64561 | 0.000153 | 0.010429 |
| *DENND1B* | 599.4 | -0.55701 | -0.67477 | 0.000829 | 0.038338 |
| *MAP4K5* | 1334.5 | 0.55277 | 0.59071 | 2.82E-07 | 5.33E-05 |
| *BCL3* | 644.3 | 0.55203 | 0.60895 | 1.56E-05 | 0.001661 |
| *LDB2* | 465.5 | -0.5518 | -0.6714 | 0.001019 | 0.044866 |
| *SMC6* | 3061.7 | -0.54865 | -0.62885 | 0.00018 | 0.01179 |
| *ZNF253* | 1646.3 | -0.54837 | -0.64474 | 0.00046 | 0.025497 |
| *NRN1* | 2783.7 | -0.54527 | -0.6447 | 0.000585 | 0.030165 |
| *SLC39A8* | 923.8 | -0.54477 | -0.65199 | 0.000815 | 0.037948 |
| *CIC* | 1809 | 0.54446 | 0.60198 | 2.46E-05 | 0.002474 |
| *SYNE3* | 1544.9 | -0.54423 | -0.60647 | 4.41E-05 | 0.004041 |
| *EVPL* | 980.6 | -0.54063 | -0.61483 | 0.000157 | 0.010539 |
| *METTL17* | 1465.2 | -0.53989 | -0.6443 | 0.000846 | 0.039027 |
| *PARVG* | 875.4 | 0.53972 | 0.62981 | 0.000445 | 0.025159 |
| *KLHL6* | 2008.1 | -0.53792 | -0.59875 | 5.01E-05 | 0.004447 |
| *ZNF772* | 554.5 | 0.53727 | 0.60117 | 7.17E-05 | 0.005934 |
| *KRT9* | 18.6 | 0.53498 | 6.65035 | 0.000955 | 0.043075 |
| *SETD5-AS1* | 404.9 | 0.53251 | 0.63055 | 0.000814 | 0.037948 |
| *IRS2* | 1062.6 | -0.52974 | -0.61918 | 0.000598 | 0.030595 |
| *TNFAIP3* | 10109.7 | 0.52792 | 0.56264 | 5.83E-07 | 9.41E-05 |
| *ASAP2* | 1227.8 | -0.52222 | -0.60451 | 0.000523 | 0.027621 |
| *HCP5* | 3037.7 | 0.52175 | 0.56108 | 3.25E-06 | 0.000412 |
| *EEF2K* | 859.8 | 0.52046 | 0.58695 | 0.000184 | 0.011958 |
| *PRKD2* | 2447.4 | 0.51983 | 0.59259 | 0.000309 | 0.018771 |
| *CTD-2368P22.1* | 306 | 0.51516 | 0.60814 | 0.001108 | 0.047917 |
| *ZNF70* | 727.1 | 0.50734 | 0.57554 | 0.000349 | 0.020819 |
| *RLTPR* | 4653.6 | -0.5058 | -0.53588 | 5.61E-07 | 9.19E-05 |
| *MYO1G* | 1804.3 | -0.50459 | -0.58325 | 0.000767 | 0.036418 |
| *TBC1D2B* | 1080.7 | -0.50443 | -0.5858 | 0.000894 | 0.040817 |
| *CORO1C* | 2814.4 | -0.50423 | -0.54246 | 7.24E-06 | 0.000851 |
| *ABHD4* | 962.1 | 0.50249 | 0.58571 | 0.001045 | 0.045721 |
| *TUBA1A* | 6080.4 | -0.49898 | -0.54946 | 8.35E-05 | 0.006682 |
| *XYLT1* | 2173.2 | 0.49304 | 0.55528 | 0.000372 | 0.021838 |
| *CAMTA2* | 666.3 | 0.488 | 0.54536 | 0.000291 | 0.017889 |
| *HERC3* | 923.4 | -0.48726 | -0.53755 | 0.000139 | 0.009751 |
| *PPP1R13B* | 1190.8 | -0.48723 | -0.53054 | 5.06E-05 | 0.004447 |
| *DARS* | 6357.7 | -0.48207 | -0.50954 | 1.17E-06 | 0.000172 |
| *HMGN5* | 399.5 | 0.48135 | 0.55117 | 0.000978 | 0.043733 |
| *RNPC3* | 603.8 | 0.47969 | 0.54177 | 0.000603 | 0.03067 |
| *NREP* | 12968.7 | -0.47495 | -0.53106 | 0.000433 | 0.024565 |
| *AHDC1* | 2906.2 | 0.47326 | 0.53736 | 0.000883 | 0.040572 |
| *STK24* | 4714.6 | -0.47268 | -0.4991 | 1.53E-06 | 0.000213 |
| *GNG2* | 3002.9 | -0.46988 | -0.5218 | 0.00035 | 0.020819 |
| *ACSS1* | 2289.9 | -0.46887 | -0.52175 | 0.000403 | 0.023211 |
| *LAT2* | 9833.8 | 0.46614 | 0.50333 | 4.94E-05 | 0.004418 |
| *MXRA7* | 1139.6 | -0.45464 | -0.49246 | 0.000101 | 0.007414 |
| *ADD3* | 2155.4 | -0.45395 | -0.48413 | 1.9E-05 | 0.00198 |
| *PRPSAP2* | 1515.8 | -0.45205 | -0.49128 | 0.000146 | 0.010068 |
| *KAZALD1* | 816.8 | 0.44361 | 0.48156 | 0.000177 | 0.011647 |
| *KIAA0930* | 1631 | -0.442 | -0.47432 | 6.48E-05 | 0.005392 |
| *CCDC117* | 2855 | -0.44158 | -0.47766 | 0.000141 | 0.009858 |
| *ZNF302* | 1339.8 | 0.43871 | 0.47895 | 0.000316 | 0.019048 |
| *MAPK13* | 1114.4 | -0.43871 | -0.48379 | 0.000588 | 0.030165 |
| *HLA-A* | 14689.3 | 0.42258 | 0.45102 | 7.51E-05 | 0.006147 |
| *ZDHHC3* | 1299.3 | -0.41703 | -0.45071 | 0.000304 | 0.018507 |
| *H2AFY* | 16915.2 | -0.41676 | -0.45876 | 0.000997 | 0.044262 |
| *PCBP1-AS1* | 1351.6 | 0.41247 | 0.44775 | 0.000489 | 0.02639 |
| *SSBP2* | 6589.7 | -0.41056 | -0.4359 | 6.44E-05 | 0.005392 |
| *SMARCA2* | 4420.1 | 0.40579 | 0.43315 | 0.000145 | 0.010068 |
| *BZRAP1* | 6827.9 | 0.38932 | 0.41596 | 0.000293 | 0.017946 |
| *ETS2* | 7236.6 | 0.38757 | 0.40978 | 9.69E-05 | 0.007272 |
| *IDH2* | 9359.9 | -0.37641 | -0.40659 | 0.001075 | 0.046676 |
| *UNG* | 4993.8 | -0.32718 | -0.3456 | 0.000909 | 0.041384 |
| *STK38* | 5328.8 | -0.32423 | -0.33965 | 0.000391 | 0.022703 |

**Supplementary table 4. Enrichment of differentially expressed genes after *E/R* abrogation.**  The first column shows the altered biological processes, following by expected value, fold enrichment, raw P value and the False Discovery Rate (FDR).

| **GO biological process complete** | **Expected** | **Fold Enrichment** | **raw P value** | **FDR** |
| --- | --- | --- | --- | --- |
| T cell receptor V(D)J recombination | .08 | 39.62 | 1.74E-04 | 3.58E-02 |
| 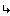somatic recombination of T cell receptor gene segments | .08 | 39.62 | 1.74E-04 | 3.53E-02 |
| 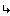immune system development | 9.63 | 2.39 | 1.98E-04 | 3.77E-02 |
| 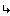system development | 66.14 | 1.57 | 9.68E-07 | 1.91E-03 |
| 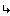[anatomical structure development](http://amigo.geneontology.org/amigo/term/GO:0048856) | 80.91 | 1.45 | 9.60E-06 | 5.42E-03 |
| 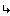[developmental process](http://amigo.geneontology.org/amigo/term/GO:0032502) | 86.12 | 1.42 | 1.45E-05 | 7.41E-03 |
| 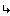[multicellular organism development](http://amigo.geneontology.org/amigo/term/GO:0007275) | 75.17 | 1.49 | 4.10E-06 | 3.42E-03 |
| 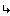[multicellular organismal process](http://amigo.geneontology.org/amigo/term/GO:0032501) | 104.29 | 1.39 | 2.50E-06 | 2.64E-03 |
| 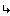[immune system process](http://amigo.geneontology.org/amigo/term/GO:0002376) | 41.50 | 1.59 | 1.53E-04 | 3.35E-02 |
| 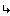[somatic diversification of T cell receptor genes](http://amigo.geneontology.org/amigo/term/GO:0002568) | .08 | 39.62 | 1.74E-04 | 3.63E-02 |
| 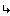[leukocyte differentiation](http://amigo.geneontology.org/amigo/term/GO:0002521) | 4.92 | 3.05 | 1.97E-04 | 3.79E-02 |
| 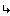[cell differentiation](http://amigo.geneontology.org/amigo/term/GO:0030154) | 55.16 | 1.50 | 1.01E-04 | 2.54E-02 |
| 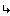[cellular developmental process](http://amigo.geneontology.org/amigo/term/GO:0048869) | 56.55 | 1.52 | 4.67E-05 | 1.64E-02 |
| 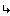[lymphocyte activation](http://amigo.geneontology.org/amigo/term/GO:0046649) | 5.63 | 2.84 | 2.60E-04 | 4.43E-02 |
| 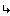[cell activation](http://amigo.geneontology.org/amigo/term/GO:0001775) | 15.87 | 2.08 | 9.11E-05 | 2.36E-02 |
| [germinal center formation](http://amigo.geneontology.org/amigo/term/GO:0002467) | .12 | 33.01 | 2.19E-05 | 1.05E-02 |
| 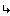[response to stimulus](http://amigo.geneontology.org/amigo/term/GO:0050896) | 126.45 | 1.27 | 1.70E-04 | 3.63E-02 |
| 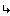[anatomical structure morphogenesis](http://amigo.geneontology.org/amigo/term/GO:0009653) | 31.84 | 1.73 | 7.23E-05 | 2.16E-02 |
| [negative regulation of necroptotic process](http://amigo.geneontology.org/amigo/term/GO:0060546) | .18 | 22.01 | 7.68E-05 | 2.21E-02 |
| 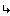[negative regulation of programmed necrotic cell death](http://amigo.geneontology.org/amigo/term/GO:0062099) | .18 | 22.01 | 7.68E-05 | 2.25E-02 |
| 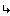[regulation of programmed necrotic cell death](http://amigo.geneontology.org/amigo/term/GO:0062098) | .27 | 14.67 | 2.88E-04 | 4.79E-02 |
| [regulation of cell death](http://amigo.geneontology.org/amigo/term/GO:0010941) | 25.40 | 2.01 | 2.34E-06 | 2.65E-03 |
| [regulation of cellular process](http://amigo.geneontology.org/amigo/term/GO:0050794) | 165.15 | 1.26 | 1.78E-06 | 2.35E-03 |
| [regulation of biological process](http://amigo.geneontology.org/amigo/term/GO:0050789) | 176.01 | 1.23 | 6.44E-06 | 4.43E-03 |
| [biological regulation](http://amigo.geneontology.org/amigo/term/GO:0065007) | 186.40 | 1.26 | 3.97E-08 | 6.29E-04 |
| [regulation of programmed cell death](http://amigo.geneontology.org/amigo/term/GO:0043067) | 23.52 | 1.87 | 8.88E-05 | 2.34E-02 |
| [negative regulation of necrotic cell death](http://amigo.geneontology.org/amigo/term/GO:0060547) | .24 | 16.51 | 1.95E-04 | 3.81E-02 |
| [negative regulation of cell death](http://amigo.geneontology.org/amigo/term/GO:0060548) | 15.02 | 2.33 | 7.20E-06 | 4.75E-03 |
| [negative regulation of cellular process](http://amigo.geneontology.org/amigo/term/GO:0048523) | 70.40 | 1.59 | 1.36E-07 | 7.16E-04 |
| [negative regulation of biological process](http://amigo.geneontology.org/amigo/term/GO:0048519) | 79.08 | 1.53 | 2.93E-07 | 1.16E-03 |
| [negative regulation of programmed cell death](http://amigo.geneontology.org/amigo/term/GO:0043069) | 13.74 | 2.26 | 4.02E-05 | 1.51E-02 |
| [regulation of necroptotic process](http://amigo.geneontology.org/amigo/term/GO:0060544) | .26 | 15.54 | 2.38E-04 | 4.23E-02 |
| [I-kappaB kinase/NF-kappaB signaling](http://amigo.geneontology.org/amigo/term/GO:0007249) | 1.05 | 6.70 | 1.40E-04 | 3.20E-02 |
| [intracellular signal transduction](http://amigo.geneontology.org/amigo/term/GO:0035556) | 25.41 | 1.89 | 2.69E-05 | 1.15E-02 |
| [signal transduction](http://amigo.geneontology.org/amigo/term/GO:0007165) | 75.12 | 1.44 | 3.56E-05 | 1.37E-02 |
| [cellular response to stimulus](http://amigo.geneontology.org/amigo/term/GO:0051716) | 99.34 | 1.33 | 1.51E-04 | 3.37E-02 |
| [signaling](http://amigo.geneontology.org/amigo/term/GO:0023052) | 80.17 | 1.47 | 3.51E-06 | 3.08E-03 |
| [cell communication](http://amigo.geneontology.org/amigo/term/GO:0007154) | 81.80 | 1.42 | 2.58E-05 | 1.13E-02 |
| [regulation of interleukin-6 production](http://amigo.geneontology.org/amigo/term/GO:0032675) | 2.01 | 4.47 | 2.88E-04 | 4.75E-02 |
| [regulation of multicellular organismal process](http://amigo.geneontology.org/amigo/term/GO:0051239) | 46.82 | 1.71 | 1.47E-06 | 2.11E-03 |
| [cellular calcium ion homeostasis](http://amigo.geneontology.org/amigo/term/GO:0006874) | 6.47 | 3.09 | 1.43E-05 | 7.53E-03 |
| [calcium ion homeostasis](http://amigo.geneontology.org/amigo/term/GO:0055074) | 6.69 | 2.99 | 2.29E-05 | 1.07E-02 |
| [metal ion homeostasis](http://amigo.geneontology.org/amigo/term/GO:0055065) | 9.42 | 2.55 | 4.26E-05 | 1.57E-02 |
| [cation homeostasis](http://amigo.geneontology.org/amigo/term/GO:0055080) | 10.60 | 2.26 | 2.57E-04 | 4.41E-02 |
| [ion homeostasis](http://amigo.geneontology.org/amigo/term/GO:0050801) | 11.80 | 2.29 | 8.73E-05 | 2.38E-02 |
| [homeostatic process](http://amigo.geneontology.org/amigo/term/GO:0042592) | 24.67 | 1.82 | 1.28E-04 | 3.11E-02 |
| [regulation of biological quality](http://amigo.geneontology.org/amigo/term/GO:0065008) | 60.69 | 1.60 | 1.30E-06 | 2.28E-03 |
| [inorganic ion homeostasis](http://amigo.geneontology.org/amigo/term/GO:0098771) | 10.78 | 2.32 | 1.38E-04 | 3.27E-02 |
| [divalent inorganic cation homeostasis](http://amigo.geneontology.org/amigo/term/GO:0072507) | 7.30 | 2.88 | 2.44E-05 | 1.10E-02 |
| [cellular divalent inorganic cation homeostasis](http://amigo.geneontology.org/amigo/term/GO:0072503) | 6.95 | 3.02 | 1.22E-05 | 6.66E-03 |
| [cellular ion homeostasis](http://amigo.geneontology.org/amigo/term/GO:0006873) | 9.66 | 2.38 | 2.04E-04 | 3.84E-02 |
| [cellular metal ion homeostasis](http://amigo.geneontology.org/amigo/term/GO:0006875) | 8.33 | 2.64 | 5.37E-05 | 1.67E-02 |
| [regulation of inflammatory response](http://amigo.geneontology.org/amigo/term/GO:0050727) | 4.98 | 3.01 | 2.23E-04 | 4.05E-02 |
| [regulation of response to stimulus](http://amigo.geneontology.org/amigo/term/GO:0048583) | 66.46 | 1.53 | 4.10E-06 | 3.25E-03 |
| [regulation of response to external stimulus](http://amigo.geneontology.org/amigo/term/GO:0032101) | 11.53 | 2.43 | 2.79E-05 | 1.16E-02 |
| [regulation of lymphocyte activation](http://amigo.geneontology.org/amigo/term/GO:0051249) | 7.72 | 2.72 | 5.35E-05 | 1.69E-02 |
| [regulation of leukocyte activation](http://amigo.geneontology.org/amigo/term/GO:0002694) | 8.97 | 2.68 | 2.00E-05 | 9.86E-03 |
| [regulation of immune system process](http://amigo.geneontology.org/amigo/term/GO:0002682) | 24.67 | 1.90 | 3.23E-05 | 1.28E-02 |
| [regulation of cell activation](http://amigo.geneontology.org/amigo/term/GO:0050865) | 9.53 | 2.73 | 6.44E-06 | 4.63E-03 |
| [regulation of GTPase activity](http://amigo.geneontology.org/amigo/term/GO:0043087) | 7.21 | 2.64 | 1.78E-04 | 3.56E-02 |
| [regulation of hydrolase activity](http://amigo.geneontology.org/amigo/term/GO:0051336) | 19.34 | 1.96 | 8.77E-05 | 2.35E-02 |
| [regulation of catalytic activity](http://amigo.geneontology.org/amigo/term/GO:0050790) | 34.97 | 1.89 | 5.47E-07 | 1.44E-03 |
| [regulation of molecular function](http://amigo.geneontology.org/amigo/term/GO:0065009) | 49.33 | 1.70 | 7.56E-07 | 1.71E-03 |
| [regulation of cell migration](http://amigo.geneontology.org/amigo/term/GO:0030334) | 12.53 | 2.55 | 2.06E-06 | 2.50E-03 |
| [regulation of cell motility](http://amigo.geneontology.org/amigo/term/GO:2000145) | 13.45 | 2.38 | 9.19E-06 | 5.38E-03 |
| [regulation of cellular component movement](http://amigo.geneontology.org/amigo/term/GO:0051270) | 14.68 | 2.45 | 1.32E-06 | 2.09E-03 |
| [regulation of localization](http://amigo.geneontology.org/amigo/term/GO:0032879) | 40.92 | 1.88 | 4.98E-08 | 3.94E-04 |
| [regulation of locomotion](http://amigo.geneontology.org/amigo/term/GO:0040012) | 14.59 | 2.19 | 4.33E-05 | 1.56E-02 |
| [actin filament-based process](http://amigo.geneontology.org/amigo/term/GO:0030029) | 8.42 | 2.49 | 1.71E-04 | 3.60E-02 |
| [regulation of cellular protein localization](http://amigo.geneontology.org/amigo/term/GO:1903827) | 8.04 | 2.49 | 2.52E-04 | 4.37E-02 |
| [regulation of protein localization](http://amigo.geneontology.org/amigo/term/GO:0032880) | 15.13 | 2.11 | 9.64E-05 | 2.46E-02 |
| [neuron projection development](http://amigo.geneontology.org/amigo/term/GO:0031175) | 9.98 | 2.40 | 1.30E-04 | 3.12E-02 |
| [cell projection organization](http://amigo.geneontology.org/amigo/term/GO:0030030) | 16.99 | 2.06 | 8.02E-05 | 2.26E-02 |
| [cellular component organization](http://amigo.geneontology.org/amigo/term/GO:0016043) | 84.69 | 1.36 | 2.08E-04 | 3.86E-02 |
| [neuron development](http://amigo.geneontology.org/amigo/term/GO:0048666) | 12.18 | 2.22 | 1.78E-04 | 3.52E-02 |
| [nervous system development](http://amigo.geneontology.org/amigo/term/GO:0007399) | 35.20 | 1.70 | 4.70E-05 | 1.62E-02 |
| [cell morphogenesis](http://amigo.geneontology.org/amigo/term/GO:0000902) | 10.56 | 2.37 | 1.07E-04 | 2.65E-02 |
| [positive regulation of immune response](http://amigo.geneontology.org/amigo/term/GO:0050778) | 12.77 | 2.19 | 1.46E-04 | 3.30E-02 |
| [positive regulation of immune system process](http://amigo.geneontology.org/amigo/term/GO:0002684) | 17.11 | 2.22 | 8.26E-06 | 5.02E-03 |
| [positive regulation of biological process](http://amigo.geneontology.org/amigo/term/GO:0048518) | 92.42 | 1.42 | 5.11E-06 | 3.85E-03 |
| [positive regulation of response to stimulus](http://amigo.geneontology.org/amigo/term/GO:0048584) | 36.33 | 1.76 | 8.16E-06 | 5.16E-03 |
| [regulation of immune response](http://amigo.geneontology.org/amigo/term/GO:0050776) | 17.01 | 2.06 | 8.09E-05 | 2.24E-02 |
| [negative regulation of apoptotic process](http://amigo.geneontology.org/amigo/term/GO:0043066) | 13.48 | 2.15 | 1.39E-04 | 3.24E-02 |
| [positive regulation of developmental process](http://amigo.geneontology.org/amigo/term/GO:0051094) | 20.46 | 2.15 | 2.91E-06 | 2.87E-03 |
| [regulation of developmental process](http://amigo.geneontology.org/amigo/term/GO:0050793) | 38.64 | 1.66 | 6.41E-05 | 1.95E-02 |
| [positive regulation of multicellular organismal process](http://amigo.geneontology.org/amigo/term/GO:0051240) | 26.05 | 2.07 | 5.37E-07 | 1.70E-03 |
| [cytoskeleton organization](http://amigo.geneontology.org/amigo/term/GO:0007010) | 16.12 | 2.05 | 1.54E-04 | 3.35E-02 |
| [negative regulation of response to stimulus](http://amigo.geneontology.org/amigo/term/GO:0048585) | 24.14 | 2.03 | 3.45E-06 | 3.21E-03 |
| [regulation of anatomical structure morphogenesis](http://amigo.geneontology.org/amigo/term/GO:0022603) | 16.05 | 1.99 | 2.70E-04 | 4.54E-02 |
| [positive regulation of molecular function](http://amigo.geneontology.org/amigo/term/GO:0044093) | 26.79 | 1.75 | 2.29E-04 | 4.11E-02 |
| [regulation of signal transduction](http://amigo.geneontology.org/amigo/term/GO:0009966) | 49.65 | 1.51 | 2.44E-04 | 4.29E-02 |
| [regulation of cell communication](http://amigo.geneontology.org/amigo/term/GO:0010646) | 55.04 | 1.53 | 5.32E-05 | 1.72E-02 |
| [regulation of signaling](http://amigo.geneontology.org/amigo/term/GO:0023051) | 55.65 | 1.55 | 2.91E-05 | 1.18E-02 |
| [positive regulation of cellular process](http://amigo.geneontology.org/amigo/term/GO:0048522) | 80.91 | 1.37 | 2.11E-04 | 3.88E-02 |
| Unclassified | 48.68 | .49 | 5.25E-05 | 1.77E-02 |

**Supplementary figure 1. Edited-REH sequences.** Sanger sequencing of the *E/R* fusion region in REH cells. The REH cells expressing PX458 without sgRNA, used as a control, had a wild type sequence, while cells expressing G1 sgRNA, G2 sgRNA or G1 + G2 sgRNAs showed a mixture of sequences around the expected Cas9 cleavage point. sgRNA guide sequences are shown in red box.

**Supplementary figure 2.** **Edited sequences of single cell derived-cell line.** The pool of REH edited-cells with both guides (G1 + G2) was separated by single cell. The sequence corresponding to the *E/R* fusion region was studied by Sanger sequencing in all clones. Two clones with single-non-edited sequence and therefore, with a wild type sequence, were selected as control clones. On the other hand, three single-edited cell clones with *E/R* KO sequence were selected as KO clones (KO1, KO2 and KO3).

**Supplementary figure 3. MTT proliferation assay.** No proliferation changes were observed in E/R KO clones respected to REH cells and control clones at 72 hours by MTT assay. This experiment had 3 replicates.

**Supplementary figure 4. Measure of tumor growth**. Mean of the tumor weight for each group of mice, represented by box plot, after being sacrificed at day 48 (group 1 and 2) or day 62 day (group 3 and 4). In this image, the mean of the sizes of the tumors is represented by box plot. In each group they are compared according to the cells injected in those mice. All the mice belonging to the same group were sacrificed on the same day. Each group had 4 replicates. **P*<0.05 (unpaired *t*-test).

**Supplementary figure 5. Tumor growth and histopathological findings.** H&E representative areas of tumors from each group. Tumors were composed of monomorphic cells with round nucleus. Although mitotic figures (yellow arrows) are observed in all tumors, a higher number were observed in REH and control tumors. Macrophages are shown in KO1 clone (box in the upper left corner).
